# Repeated exposure to a conserved antigen drives dynamic clonal and epitope succession

**DOI:** 10.64898/2026.11.12.763997

**Authors:** Hiroyasu Aoki, Mikhail V. Pogorelyy, Philip Bradley, Cecilia Adragna, Stefan A. Schattgen, Anastasia A. Minervina, Koshlan Mayer-Blackwell, Kasi Vegesana, Zachary B. Scott, Velmurugan Balaraman, E. Kaitlynn Allen, Masahiro Kitabatake, Noriko Ouji- Sageshima, Atsushi Hara, Peng Xu, Chihiro Motozono, Kouji Matsushima, Federica Sallusto, Antonio Lanzavecchia, Satoshi Ueha, Toshihiro Ito, Paul G. Thomas

**Author notes:** Corresponding author. (S.U.), (P.G.T). Currently employed by ImmunoGeneTeqs, Inc (Peng Xu).

## Abstract

Defining the dynamics of T cell memory formation with epitope-level resolution can inform vaccine design. Longitudinal T cell receptor (TCR) sequencing after COVID-19 mRNA vaccination has enabled detailed tracking of T cell responses; however, connecting expanded clonotypes following antigen exposure to their cognate epitopes remains a challenge due to limited curated paired α/βTCR information. To fill this gap, we used TIRTL-seq to identify paired α/βTCRs from CD8^+^ T cell lineages associated with mRNA immunization in serial samples across 38 healthy donors over multiple rounds of vaccinations. Vaccine-associated paired α/βTCRs, identified based on their longitudinal kinetics, were grouped according to sequence similarity to infer their shared epitope specificities. We assigned epitope specificity to 1,109 clonotypes, recognizing 15 SARS-CoV-2 spike targets, identifying multiple previously undescribed responses, and confirming putative specificities experimentally. This approach uncovered a striking pattern of clonal succession in anti-spike T cell memory lineages resulting from repeated vaccination, characterized by a transition from low-avidity clones recognizing conserved coronavirus epitopes to high-avidity clones targeting epitopes unique to SARS-CoV-2. These results highlight the surprisingly dynamic responses underlying homologous boosting, suggest avenues for improving vaccine strategies, and pave the way for repertoire-based monitoring of antigen-specific T cell immunity.

## Introduction

mRNA vaccines using the spike glycoprotein of severe acute respiratory syndrome coronavirus 2 (SARS-CoV-2) play a significant role in reducing the burden of coronavirus disease 2019 (COVID-19)^1,2^. One reason for the success of mRNA vaccines is their ability to induce robust cellular immune memory mediated by CD4^+^ and CD8^+^ T cells, as spike-specific T cells are more resilient to mutations occurring in SARS-CoV-2 variants of concern (VOCs) than antibody-mediated humoral immunity^3–5^. Thus, an in-depth understanding of vaccine-induced T cell responses is critical for assessing the efficacy and durability of mRNA vaccines against an evolving SARS-CoV-2 virus.

T cells comprise diverse clonal populations with distinct antigen specificities, each defined by a unique T cell receptor (TCR) generated through V(D)J recombination. High-throughput TCR-sequencing (TCR-seq) performed at critical time points prior to exposure and during the acute and/or convalescent phase enables characterization of vaccine-induced T cell responses based on clonal expansion in an antigen-agnostic manner, in contrast to pooled peptide stimulation^6,7^ or peptide HLA (pHLA) multimer staining^8,9^, which often requires prior knowledge of T cell epitopes.

We established a cohort of 38 healthcare workers in Japan (Nara Medical University: NMUCoV cohort) who were naive to SARS-CoV-2 at enrollment and subsequently received three or more mRNA vaccines for up to three years and sampled these donors at uniform time points to capture both acute and convalescent phases of vaccine-induced T cell responses^10,11^. By performing longitudinal TCR-seq on 38 donors of this cohort, we previously reported that the kinetics of vaccine-induced CD8^+^ T cells differ across clonotypes^12^. Specifically, some putatively SARS-CoV-2-specific clonotypes expand rapidly to the priming dose but show limited expansion in response to later boosters, and they are apparently succeeded by other clonotypes that persist as stable memory ^11^. However, the epitope specificities of the majority of vaccine-induced TCRs remain unknown, limiting immunological or mechanistic interpretations of these findings.

Since both the α and β chains comprising the TCR are essential for antigen recognition, obtaining sequences of paired α/βTCRs is necessary to experimentally identify their target epitope and quantify their functional avidity by in vitro reconstitution^13^. In this study, we sought to produce a comprehensive profile of repeated vaccine-induced responses with epitope-level resolution. To accomplish this, we applied TIRTL-seq (Throughput-Intensive Rapid TCR Library sequencing)^14^ on 124 PBMC samples collected from our cohort to significantly increase the number of recovered paired α/βTCRs compared to other methods^15,16^, allowing us to expand the landscape of antigen specificity of vaccine-induced TCRs with annotated clonal dynamics. By integrating TIRTL-seq data with longitudinal TCRβ-seq results^11,12^, we identified 5,681 paired αβTCRs of CD8^+^ T cell clones associated with vaccination and grouped them based on sequence similarity to infer their shared epitope specificities. Using donors’ HLA allele information and published databases of spike-specific TCRs, together with novel AlphaFold3-based TCR-pHLA structural modeling^17,18^, we assigned 15 SARS-CoV-2 spike epitope specificities to 1,109 TCRs, including 11 epitopes for which no cognate TCR had previously been determined, with confirmatory experimental validation. These results uncovered a surprisingly dynamic clonal succession in anti-spike T cell memory induced by repeated vaccination with an identical antigen, characterized by a transition from clones recognizing conserved coronavirus epitopes to those targeting epitopes unique to SARS-CoV-2.

## Results

### Classifying CD8 clonal response patterns induced by repeat mRNA vaccination

We longitudinally collected PBMC samples from a cohort of 38 healthcare workers in Japan (“NMUCoV” cohort) who received three or more SARS-CoV-2 mRNA vaccinations (donors’ metadata and sample availability are summarized in Supplementary Table 1). We have previously performed bulk TCRβ-seq across 16 time points over three years, which provided the response kinetics of TCRβ clones following repeated vaccinations (Fig. 1a)^11,12^. In this study, we performed TIRTL-seq on 124 PBMC samples at 1-week or 1-month after each vaccination to obtain paired α/βTCRs of CD8^+^ T cell clones induced by repeated mRNA vaccination (Fig. 1a). Collectively, these data reflect the trajectories of 19,112,208 independent clonal lineages.

**Fig. 1.**
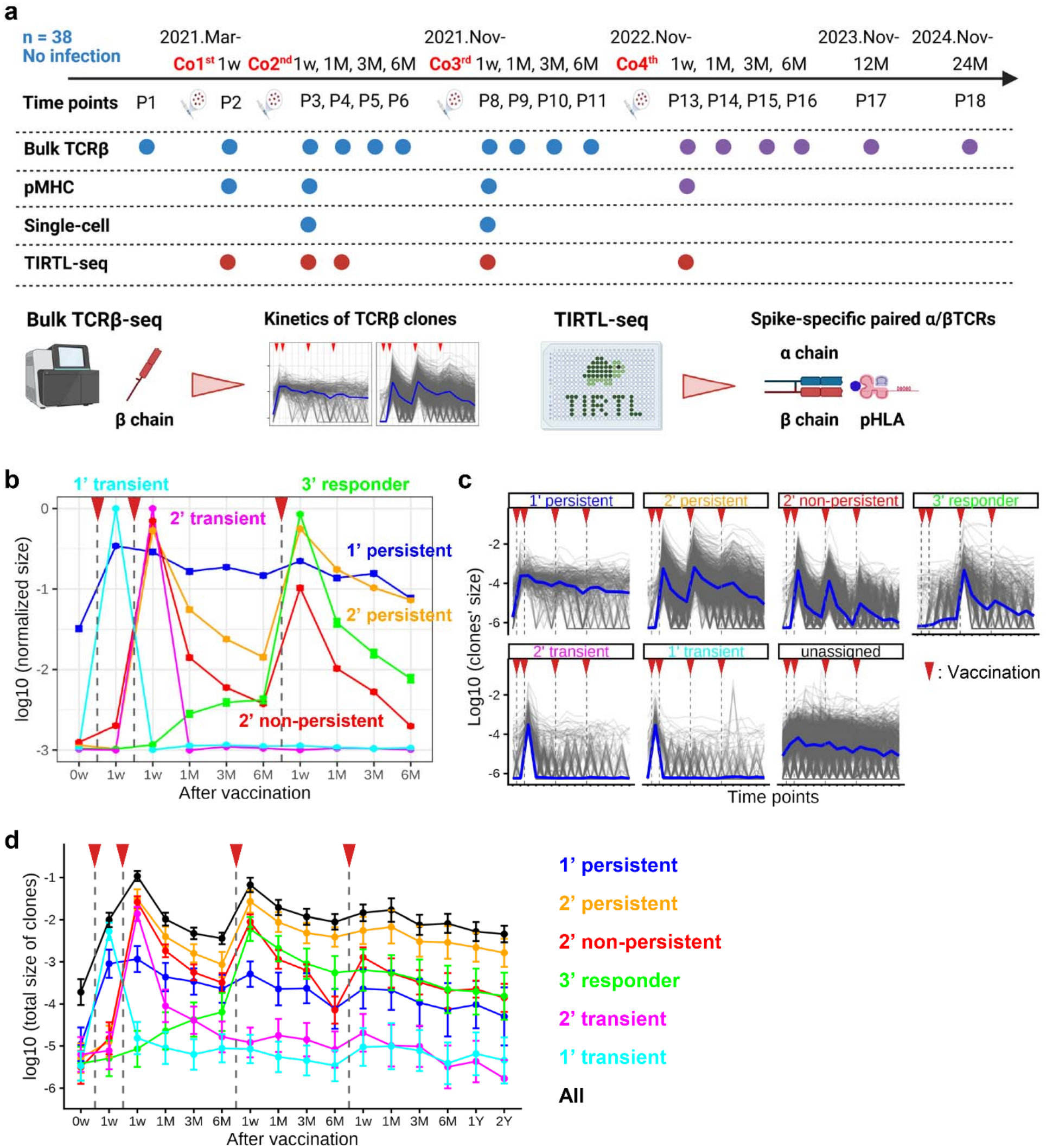
Classifying vaccine-induced CD8 responses into distinct response patterns. (**a**) Schematic of longitudinal sample collection. Circles indicate the time points at which bulk TCRβ-seq, spike pHLA tetramer TCR-seq, and TIRTL-seq were performed. Circle colors indicate the study in which the data were first reported (blue; Aoki et al.^12^, purple; Xu et al.^11^, red; this study). (**b**) Response patterns of vaccine-induced clones identified by PCA and k-means clustering on their longitudinal kinetics (Extended Data Fig. 1a-c). For each clone, the size at each time point was normalized to its maximum value across P1–P11 and then transformed to a log_10_ scale. (**c**) Kinetics of individual CD8^+^ clones classified into one of the six response patterns or left unassigned. Blue lines represent the average of log_10_-transformed clone frequency in each response pattern. (**d**) Decomposing vaccine-induced CD8^+^ T cell responses into the six response patterns. For each donor, the total size of vaccine-induced clones was calculated within each response group and for all groups combined (“All”). The mean and 95% confidence interval (CI) of log_10_-transformed values across donors are shown (n=38). Red triangles indicate the timing of vaccination. (**a,c**) Created using BioRender.com.

First, using our longitudinal bulk TCRβ-seq dataset, we identified vaccine-induced T cell clones that expanded after any of the four vaccinations relative to their pre-vaccination levels and classified them based on their longitudinal kinetics in an unsupervised manner (Extended Data Fig. 1a). Through this analysis, we identified 8,396 vaccine-induced CD8^+^ T cell clones (Fig. 1b,c, Extended Data Fig. 1b-f, and Supplementary Table 2; see *Methods*) and further decomposed the overall vaccine-induced CD8^+^ T cell response, measured as the total frequency of vaccine-induced clones per donor (Fig. 1d, black line), into six response patterns (Fig. 1d, colored lines). 5,954 clones peaked after the 2^nd^ dose and were classified into three patterns: those that re-expanded after the 3^rd^ dose and then persisted (*2’ persistent*: 1,461 clones), those that re-expanded but did not persist (*2’ non-persistent*: 1,605 clones), and those that did not re-expand (*2’ transient*: 2,888 clones). In addition, 1,827 clones that peaked after the 1^st^ dose were divided into two patterns: persistent (*1’ persistent*: 295 clones) or transient (*1’ transient*: 1,532 clones). Finally, 615 clones peaked after the 3^rd^ dose, representing the sixth response pattern (*3’ responder*) (Extended Data Fig. 1g,h). The average vaccinee showed that ∼10% of all circulating CD8 cells were vaccine-induced after the second shot, with subsequent vaccine boosts reaching estimated magnitudes of ∼6% of all CD8 T cells as vaccine-responsive. These results suggested that repeated mRNA vaccination progressively skews the CD8^+^ T cell responses towards a subset of persistent vaccine-induced clonotypes.

### Persistent vaccine-induced CD8^+^ T cell clones are characterized by slower decay kinetics and stronger activation phenotypes

To characterize CD8^+^ clones recruited as persistent memory after repetitive mRNA vaccination, we compared the longitudinal kinetics and gene expression profiles between *2’ persistent* and *2’ non-persistent* vaccine-induced clones, both of which peaked after the second dose. First, we examined when the longitudinal kinetics of persistent and non-persistent clones began to diverge. We used a generalized linear model to classify the two groups based on clone size at each time point and sequentially added time points used in the model, starting from the earliest to the latest (Fig. 2a). The model performance, measured by ROC-AUC, increased from 0.56 to 0.73 when incorporating the data from 1 month after the 2^nd^ dose (Fig. 2b,c), indicating that the clone size at this time point is an important discriminator between *2’ persistent* and *2’ non-persistent* clones. To confirm this, we compared the size and fold changes at each time point between the two groups. There was only a small difference in average clone sizes between persistent and non-persistent clones 1 week after vaccination (0.056% vs 0.044%); however, by 1 month after vaccination, the persistent clones had contracted less than non-persistent clones (0.0050% vs 0.0007%; Fig. 2d and Extended Data Fig. 2a). Whereas non-persistent clones decreased on average by 98% from peak size within 1 month, persistent clones decreased by only 90% (Fig. 2e and Extended Data Fig. 2b). Given the absence of time points between 1 week and 1 month after the 2^nd^ dose, these results suggest that clones exhibiting either i) slower contraction or ii) a delayed, later-peaking response after initial prime-boost mRNA vaccinations are more likely to survive as long-term memory.

**Fig. 2.**
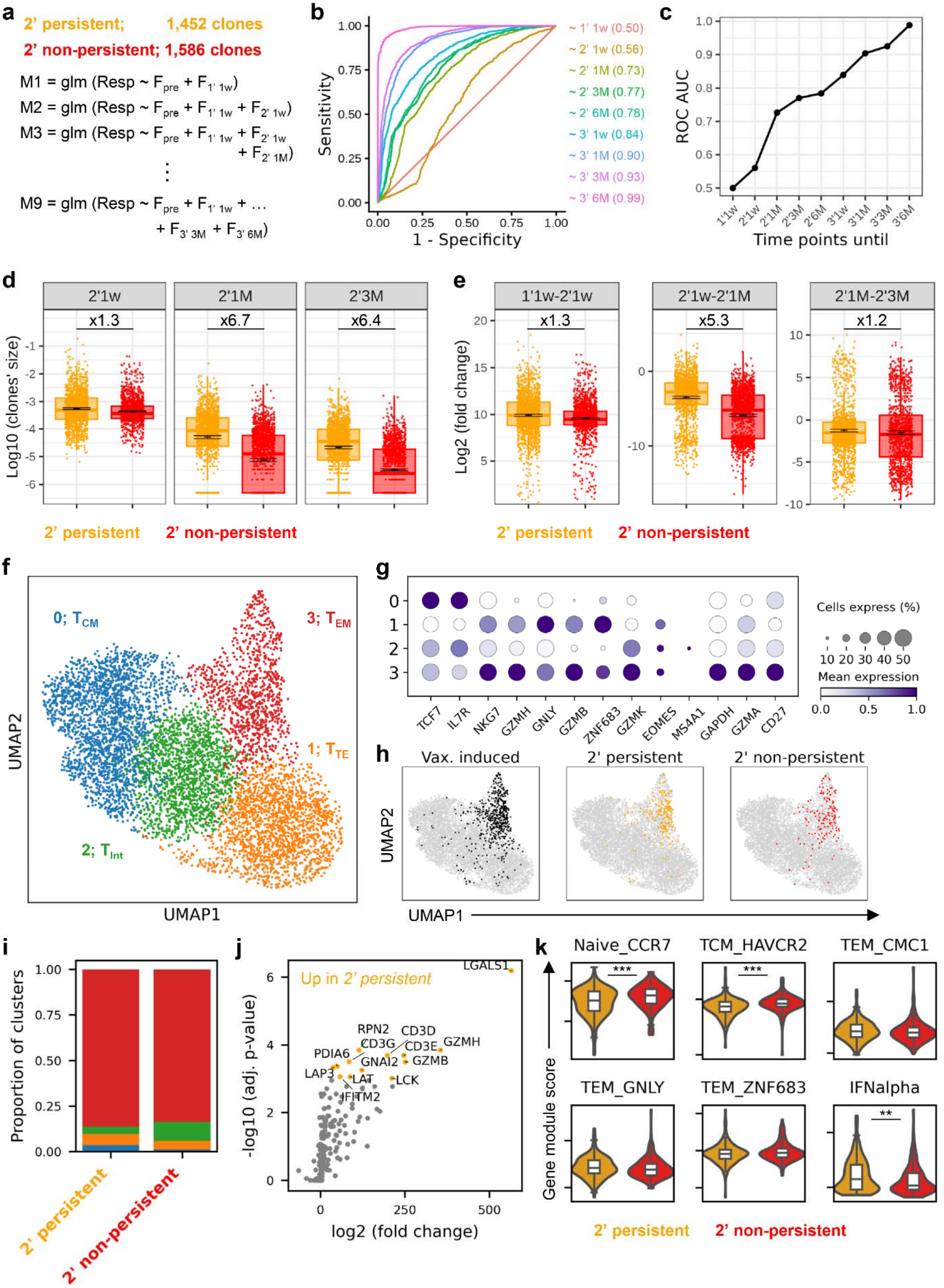
Characterizing persistent CD8^+^ T cell clones induced by vaccination. (**a**) Overview of generalized linear models to classify *2’ persistent* and *2’ non-persistent* vaccine-induced clones based on their longitudinal kinetics. F_XX_ means the frequency of a clone at the time point XX. (**b**) ROCs of the models. (**c**) ROC-AUC values of the different models. The last time point used in each model was indicated in the legend (b) or plotted on x-axis (c). (**d**) Size of individual *2’ persistent* and *2’ non-persistent* clones at 1 week (left), 1 month (middle), and 3 months (right) after the 2^nd^ dose. (**e**) Fold changes in the size of individual *2’ persistent* and *2’ non-persistent* clones between 1 week after the 1^st^ and 2^nd^ doses (left), 1 week and 1 month after the 2^nd^ dose (middle), and 1 month and 3 months after the 2^nd^ dose (right). Horizontal lines and bars represent the mean and 95% CI of log_10_-transformed values, and box plots represent the median and the interquartile range. The fold differences in the mean values between the two groups are also shown. *2’ persistent*; n=1,452 clones. *2’ non-persistent*; n=1,586 clones. (**f**) Clonotype-level UMAP of gene expression. Each dot represents a CD8^+^ T cell clone, with gene expressions averaged across cells within that clone. (**g**) Dot plot showing mRNA expression of marker genes in the clusters of T cell clones. Dot size represents the percentage of clones in each cluster expressing the gene, and dot color indicates the average gene expression level per clone. (**h**) Clonotype-level UMAP plot depicting vaccine-induced clones (left), as well as *2’ persistent* (middle) and *2’ non-persistent* (right) clones at 1 week after the 2^nd^ dose (same as below). (**i**) Bar plot representing the cluster distribution of *2’ persistent* and *2’ non-persistent* clones. (**j**) Volcano plots of differentially expressed genes between *2’ persistent* and *2’ non-persistent* clones. Genes expressed more than 8-fold with statistical significance (p < 0.001) in *2’ persistent* clones are colored in orange. (**k**) Violin plot showing gene signature scores of *2’ persistent* and *2’ non-persistent* clones. Signature genes are curated from the marker genes of T cell clusters described in Wang et al.^20^, or Liberzon et al^21^. P values of Student’s t-test were shown. ***p<0.001; **p<0.01. (**a**-**e**) Clones from Donor 018, who lacks data at P11, were excluded. (**h**-**k**) Data reported in Aoki et al. were re-analyzed (n=7)^12^.

To assess whether contraction kinetics were linked to post-vaccine phenotypes, we compared the gene expression profiles of *2’ persistent* and *2’ non-persistent* clones by analyzing a single-cell RNA/TCR-seq dataset of peripheral blood CD8^+^ T cells from 7 donors at 1 week after either the 2^nd^ or 3^rd^ dose, obtained in our previous study (Extended Data Fig. 2c,d)^12^. We averaged gene expression across all cells within each clone and generated a UMAP plot of these CD8^+^ T cell clones with Leiden clustering (Fig. 2f,g). We found that most vaccine-induced, *2’ persistent*, and *2’ non-persistent* clones exhibited T_EM_ phenotypes at 1 week after the 2^nd^ dose (Fig. 2h,i).

Notably, at the same time, *2’ persistent* clones expressed significantly higher levels of effector molecules (*GZMB* and *GZMH*) and T cell activation-induced genes (*LGALS1*)^19^ compared with *2’ non-persistent* clones (Fig. 2j). Consistently, *2’ persistent* clones showed lower module scores of CD8^+^ naïve and central memory marker genes^20^, while they had higher scores of IFNα-induced genes (Fig. 2k)^21^. These results suggest that clones receiving stronger TCR signaling or type-I IFN stimulation during the acute response are preferentially retained as stable memory after mRNA vaccinations.

### Identifying clusters of vaccine-associated CD8^+^ clones putatively sharing a target epitope

By combining the paired α/βTCR sequences obtained by TIRTL-seq and single-cell TCR-seq with the longitudinal TCRβ-seq dataset, we identified 105,880 paired CD8^+^ T cell α/βTCRs in 38 donors, covering ∼60% of CD8^+^ T cells on average. From them, we classified 4,473 TCRs as putatively vaccine-induced based on their longitudinal kinetics (Fig. 3a). In addition, we re-analyzed the complete data set using our vaccine-induced clones as “bait” to rescue CD8^+^ TCRs that showed high sequence similarity to those of vaccine-induced clones, and thus are likely spike specific, but whose longitudinal kinetics failed to meet the criteria of “vaccine-induced” due to weaker clonal expansion (Fig. 3a and Extended Data Fig. 3a, *Method*). We confirmed the validity of this sequence neighbor-selection approach by examining the longitudinal kinetics of spike-specific pHLA tetramer^+^ T cell clones detected using bulk TCR-seq (“*spike pHLA tetramer TCR-seq*”), performed in our previous study (Extended Data Fig. 3b)^11,12^. Among tetramer^+^ clones not classified as vaccine-induced in our initial screen, those rescued by TCR sequence similarity on average showed expansion after vaccination that fell below the strict statistical thresholds we set for the unsupervised kinetic clustering analysis. In total, recruiting neighbor sequences identified approximately 1,200 additional putative SARS-CoV-2-specific CD8^+^ TCRs, totaling 5,681 CD8^+^ clones across 38 donors (listed in Supplementary Table 3).

**Fig. 3.**
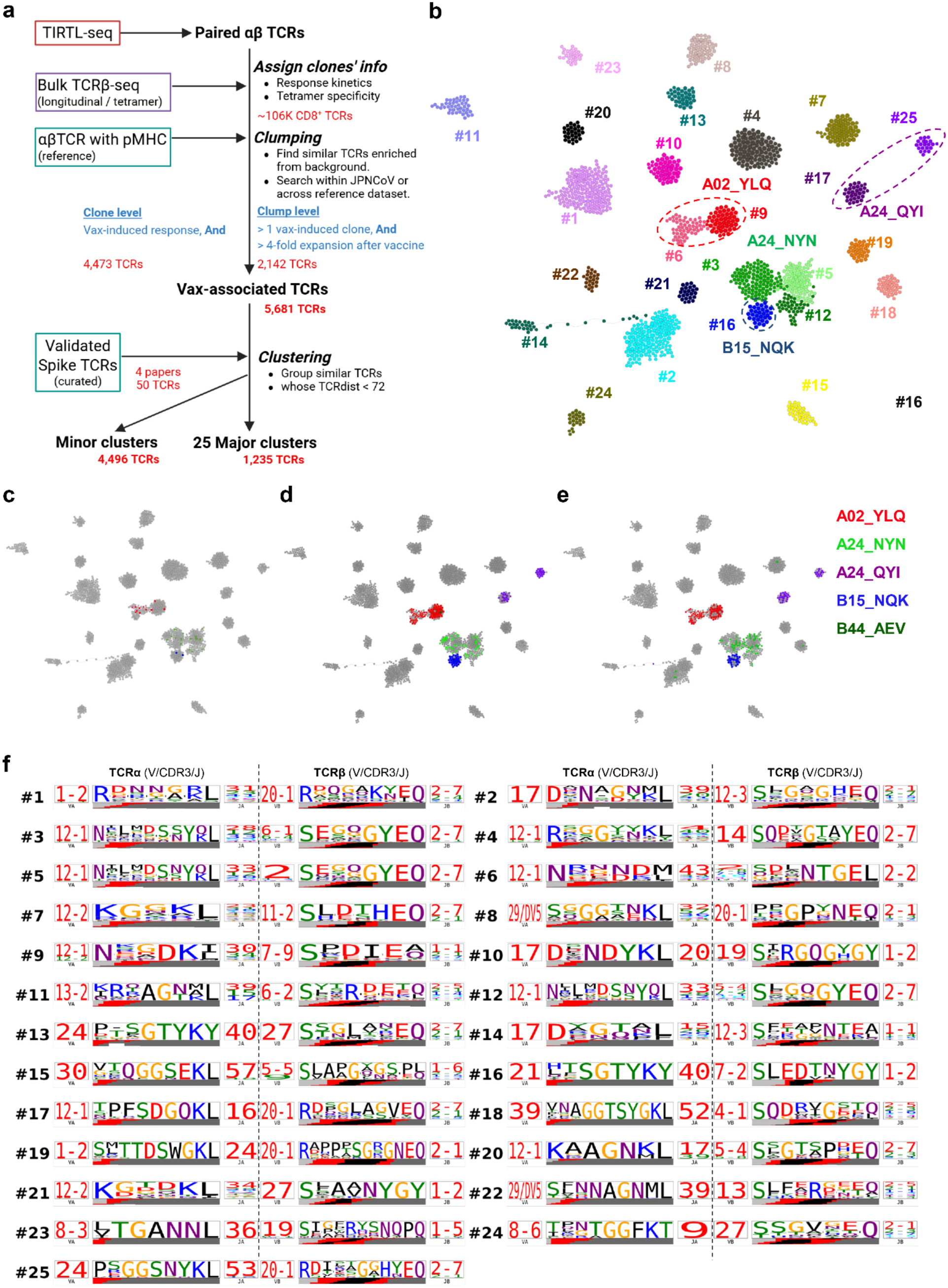
Clustering vaccine-associated CD8^+^ clones based on their α/βTCR sequences obtained by TIRTL-seq. (**a**) Workflow for identifying clusters of vaccine-induced CD8^+^ T cell clones with similar TCR sequences. (**b**) TCR network graph of 25 major clusters of vaccine-associated CD8^+^ clones. Each node represents a clone, colored by clusters, and edges connect clones with high TCR sequence similarity (tcrdist ≤ 72). (**c**-**e**) De-orphanized spike-specific TCRs, which were validated by in vitro functional assays in other studies (**c**)^24–26^, estimated by the public TCR-pHLA database (**d**) or obtained from spike pHLA tetramer TCR-seq in our previous study (**e**). (**f**) Amino acid sequence motifs (TCR logos) of paired αβTCRs for the 25 major clusters. (**a**) Created using BioRender.com.

Next, to estimate the target epitopes of these vaccine-associated clones utilizing the principle that T cells recognizing the same epitope often have similar TCR sequences^22,23^, we clustered our vaccine-induced clones based on their TCR sequence similarity. This analysis identified 25 major TCR clusters (each with at least 20 vaccine-associated TCRs shared across multiple donors), comprising 1,235 TCRs (Fig. 3a,b, and Extended Data Fig. 3c). Moreover, we annotated TCRs whose spike epitope targets were previously validated (“de-orphanized” TCRs) in our lab (Fig. 3c)^24–26^, along with TCRs matching spike-specific α/βTCRs in public databases such as VDJdb (Fig. 3d)^25–32^, and TCRs detected in our spike pHLA tetramer TCR-seq (Fig. 3e)^11,12^. This confirmed that TCRs with different specificities were well segregated into different clusters.

The TCR sequence motifs shared within each cluster, visualized as TCR logos^23^, highlight differential conservation of TCRβ versus TCRα motifs across distinct clusters (Fig. 3f). For example, clusters 9 and 16, which contain previously de-orphanized TCRs specific to A*02:01-presented S_269-277_ (A02_YLQ) and B*15:01-presented S_919-927_ (B15_NQK), respectively, show stronger conservation in their CDR3β V/D/J junction sequences compared to CDR3α, while in contrast, clusters 17 and 25 (specific to A*24:02-presented S_1208-1216_: A24_QYI) show a clearer sequence motif in their CDR3α V/J junction.

### Estimating and experimentally validating spike target epitopes of CD8^+^ TCR clusters

Among the 25 major TCR clusters, eight clusters could be directly linked to a previously characterized peptide-HLA (pHLA) epitope using published TCRs specific to these epitopes (Fig. 3c-e; Supplementary Table 4, column 4)^11,12,24–32^. Next, we tried to predict the target pHLA for the remaining vaccine-associated TCR clusters (*Methods*, in detail). First, we inferred HLA allele restriction of TCR clusters using the typing results of HLA-A and -B alleles of donors contributing to each cluster and examining their shared HLA overlap (Fig. 4a, blue box; Supplementary Table 4, column 5). This allowed us to narrow down candidate spike epitopes to those predicted to bind the inferred HLA allele using NetMHCpan-4.1^33^. Second, based on prior studies, we curated a reference set of 26,266 TCRβs associated with spike peptide reactivity or HLA-enrichment after SARS-CoV-2 exposure from two published datasets^34,35^ (Fig. 4a, green box) and mapped them to our set of vaccine-associated TCRs, allowing for one amino acid difference between the reference and query CDR3β sequences (“edit-1 neighbors”). For each cluster, we then identified the most frequently shared edit-1 neighbor and assigned its associated spike epitope or HLA allele as the putative target of that cluster. This allowed us to assign a candidate pMHC to four more clusters (e.g., B35_SSA to cluster 4 and A24_RFP to cluster 7; Supplementary Table 4, column 6).

**Fig. 4.**
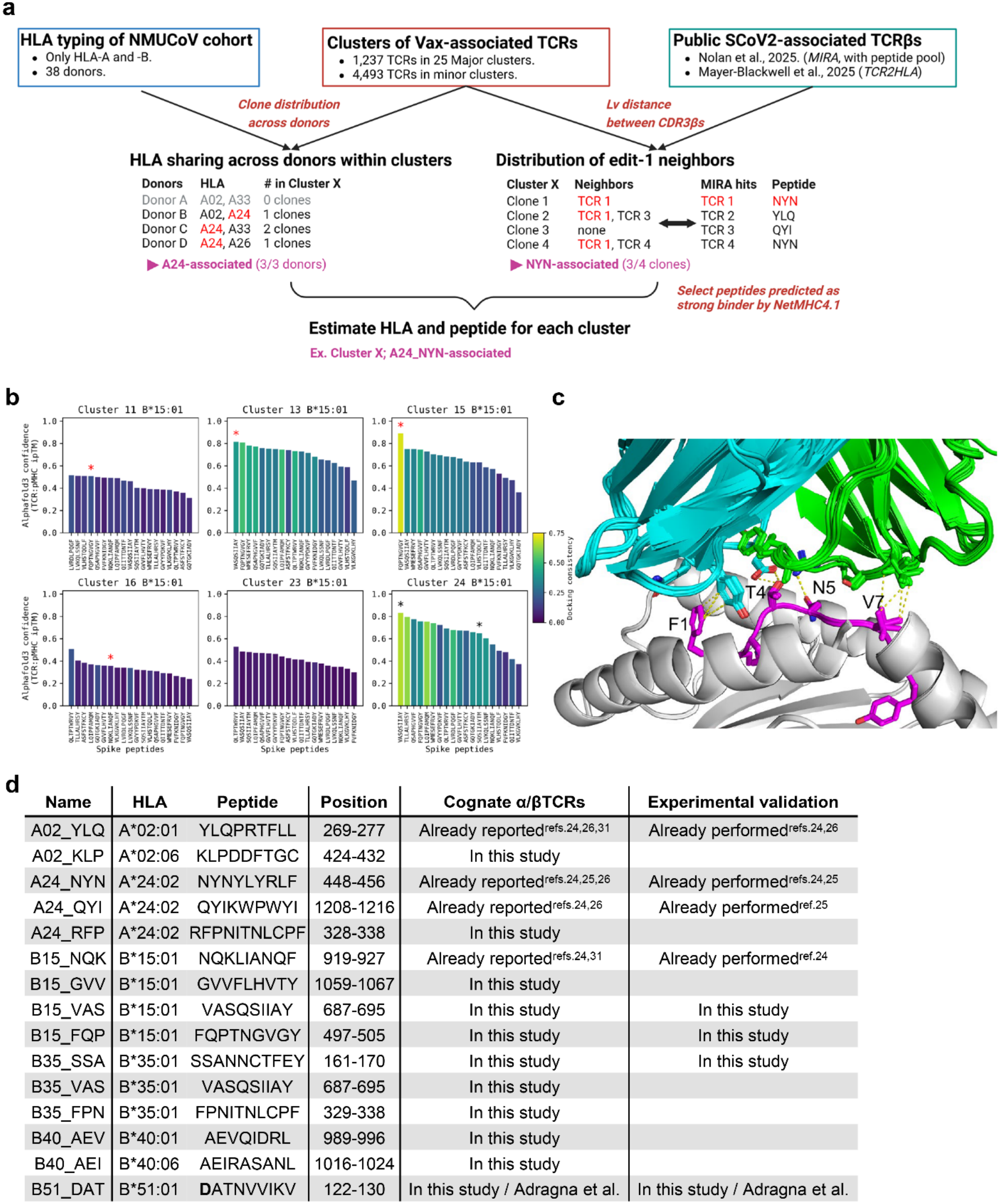
Predicting spike epitopes of CD8^+^ vaccine-associated clones by TCR clustering-based approach. (**a**) Workflow for predicting the target peptide and HLA alleles of TCR similarity clusters of vaccine-induced CD8^+^ T cell clones. Please refer to the methods section in detail. (**b**) Structure-based peptide prediction for six major TCR clusters derived from HLA-B*15:01^+^ donors. 3D models of TCR:pMHC complexes for 9-mer spike epitopes predicted to be strong binders to B*15:01 were generated using the *TCRdock* pipeline with the AlphaFold3 prediction network. AlphaFold confidence in the predicted interface, measured by the mean TCR:pMHC interchain interface-predicted template-modeling (ipTM) score, is shown as bars. Bar colors indicate the consistency of modeled TCR:pMHC docking geometries across TCRs, an additional indicator of modeling confidence. Asterisks show the estimated (in black) or experimentally confirmed (in red) target spike epitopes of the clusters (Extended Data Fig. 5). (**c**) Top 10 predicted models of TCR:pMHC complexes for TCRs of Cluster 15 with the FQP peptide presented on HLA-B*15:01. Predicted interactions between TCR CDR3 and the FQP peptide are shown as dashed yellow lines. CDR3α: cyan; CDR3β: green; HLA-B*15:01: gray; FQP peptide: magenta. (**d**) List of SARS-CoV-2 spike epitopes whose cognate α/βTCRs were identified in this study. Positions in amino acid sequence of SARS-CoV-2 spike protein (original Wuhan strain) are shown. B51_DAT epitope is produced by deamidation of position 1 Asn. (**a**) Created using BioRender.com.

In parallel, we attempted to predict the targets of vaccine-associated TCR clusters in the context of the inferred HLA allele by modeling the 3D structures of TCR-pMHC interaction using AlphaFold3^17,18^: we ran AlphaFold3 simulations for each TCR in the cluster against all 9-mer spike peptides predicted to strongly bind the associated HLA allele, and then computed the average score of model confidence as well as the similarity of the predicted TCR:pMHC docking geometries within each cluster (Fig. 4b,c, and Extended Data Fig. 4; *Methods*). This 3D structure-based prediction provided high-confidence predictions in only a few cases, including the binding of TCRs of cluster 6 to A02_YLQ (Extended Data Fig. 4). However, it strongly identified a candidate epitope for the B15-associated cluster 15, which exhibited a high binding score to the FQPTNGVGY peptide on HLA-B*15:01 (B15_FQP) with consistent docking geometry (Fig. 4b,c).

After using this TCR clustering-based approach, we could predict putative spike epitopes for 13 of 25 major TCR clusters as well as 22 minor clusters (Supplementary Tables 4 and 5). We next set out to experimentally validate the specificities of 9 clusters by transducing their TCRs into an NFAT-GFP reporter Jurkat cell line and performing a peptide co-culture assay (Extended Data Fig. 5a,b). In addition to these 9 clusters, we cloned TCRs from 4 clusters whose HLA restrictions were predicted and tested their reactivity to spike peptides predicted to strongly bind the inferred HLA allele^33^ or reported as epitopes presented on the allele (*Adragna et al., from personal communication*). From this, we found that representative TCRs from clusters 2, 10, and 14 recognize the B51_DAT epitope, a novel epitope not previously reported, while those from cluster 11 recognize the B15_FQP epitope (Extended Data Fig. 5a,b). Interestingly, the B51_DAT epitope requires a post-translational deamidation of asparagine to aspartic acid (N to D at position 1) of the vaccine-encoded peptide NATNVVIKV (S_122-130_), which was first identified by immunopeptidomics (*Adragna et al., from personal communication*). This is consistent with the fact that B51-DAT-specific clusters (2, 10, and 14) mapped to many reference TCRs with strong evidence of HLA-enrichment after SARS-CoV-2 spike exposure^34^ but failed to map to a database of TCRs recognizing spike peptides without modification^35^.

Possibly because of this unconventional pathway of peptide production, the reactivity of B51_DAT-specific TCRs varied greatly across and within TCR clusters in our in vitro co-culture assay; all three TCRs from cluster 10 and one TCR from cluster 2 showed reactivity comparable to other spike-specific TCRs, while 6 TCRs from clusters 2 and 14 only recognized the peptide at high concentration of 50 μM, and the remaining 4 representative TCRs did not become GFP^+^ even at this high peptide concentration (Extended Data Fig. 5a,b), suggesting that there may be additional peptide modifications to this epitope required for recognition in vivo.

In addition to the TCR clustering-based epitope prediction, we could infer the specificities of an additional 124 clones from sequence matches to our spike pHLA tetramer TCR-seq results or from sequence similarity to TCRs previously identified in other datasets^11,12,25–32^. In summary, from 38 donors, we could assign specificity to 1,109 vaccine-associated paired α/βTCRs, 72 of which were experimentally validated across 13 clusters. This combined approach reconfirmed TCR assignments to four prior immunodominant epitopes and allowed us to de-orphanize paired α/βTCRs for 11 additional spike epitopes, including two epitopes presented on HLA-B*15:01 (B15_VAS and B15_FQP), that lacked prior assignment of paired α/βTCRs and in vitro validation (Fig. 4d).

### Deconvoluting the vaccine-induced CD8^+^ T cell response into distinct spike epitopes

Using the list of TCR clusters of vaccine-associated clones with predicted spike target epitopes, we further deconvoluted CD8^+^ clonotype-level vaccine-induced responses into distinct spike epitope trajectories. From this, the total size of clones assigned to each spike epitope was calculated and tracked longitudinally within each donor (Fig. 5a,b). On average, these specificity-assigned clonotypes occupy 1.02% (median, ranging 0.016–8.01%) of total CD8^+^ T cells at 1 week after the 2nd dose. At the epitope level, clonotypes assigned to A02_KLP, B51_DAT, and B35_SSA accounted for more than 1% of total CD8 T cells 1 week after the 2^nd^ dose, averaged across donors with the corresponding HLA allele (median). Moreover, these plots quantify the distinct kinetics of CD8^+^ T cell responses across spike epitopes: most epitopes, including A02_YLQ, induce conventional, repetitively boosting responses to 2^nd^ and subsequent doses (categorized as *2’ persistent* or *2’ non-persistent*), whereas responses to B15_NQK and B40_AEV show an early-responder pattern (*1’ persistent*); responses to A24_NYN and A24_QYI show a mixture of these two patterns. Notably, more than half of the vaccine-associated CD8^+^ repertoire still lacks predicted spike epitopes because they did not match any of the published SARS-CoV-2-associated TCRs and are not clustered with any of the newly de-orphaned specificities due to low TCR similarity with other clones (Extended Data Fig. 6).

**Fig. 5.**
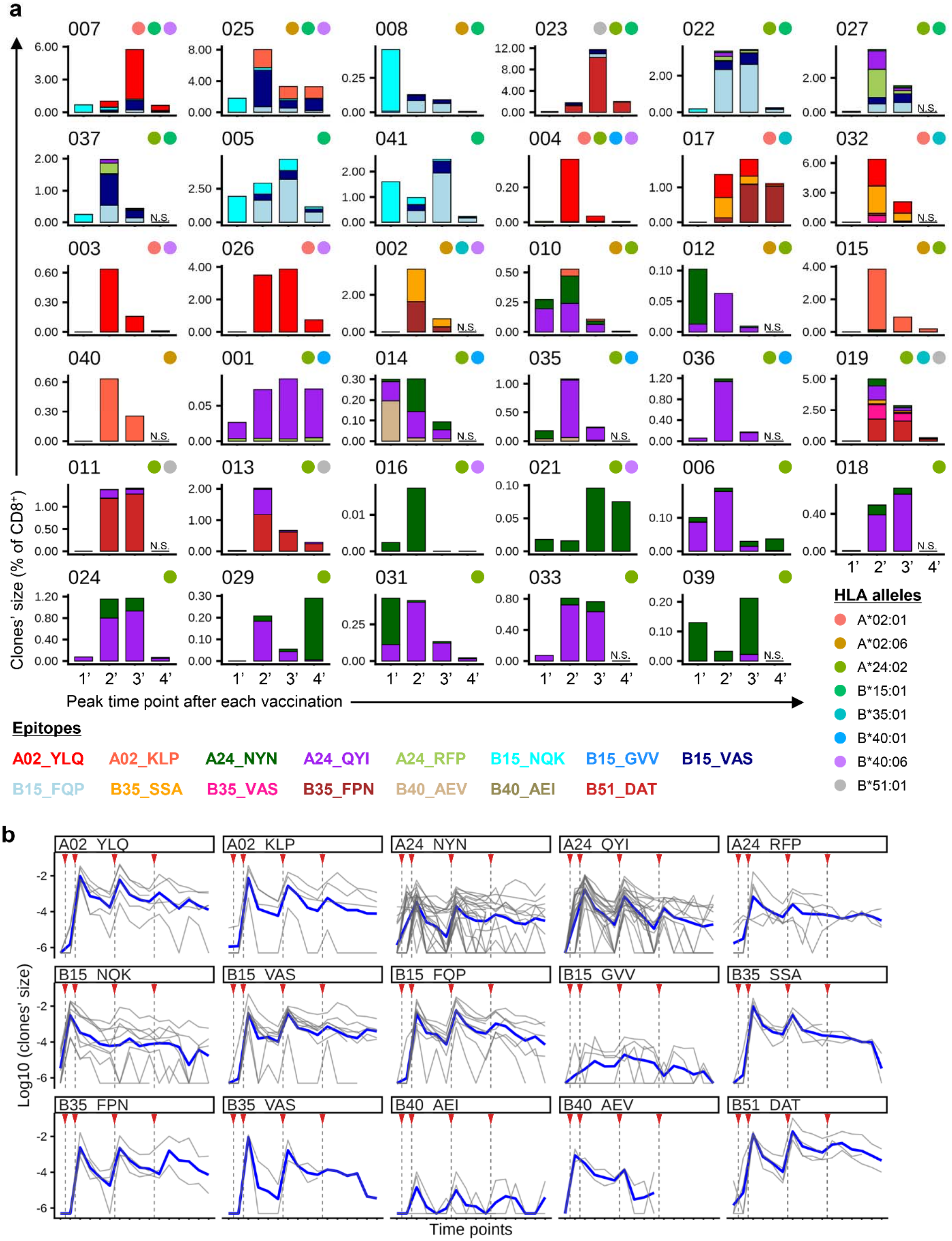
Deconvolute vaccine-induced CD8^+^ T cell responses into distinct antigen specificities. (**a**) CD8^+^ T cell responses deorphanized to individual spike epitopes within each donor. Bar height represents the total size of CD8^+^ clones deorphanized to each epitope, indicated by its color. Numbers and colored circles above the plots denote individual donors and HLA alleles they carry. Only HLA alleles that present spike epitopes predicted for TCR clusters (as summarized in Fig. 4d) are shown. Donors 030 and 038 are excluded because they lack these HLA alleles, and Donor 020 is not shown due to the absence of deorphanized CD8^+^ clones. 1’: 1 week after the 1^st^ dose. 2’: 1 week after the 2^nd^ dose. 3’: 1 week after the 3^rd^ dose. 4’: 1 month after the 4^th^ dose. N.S.: Not sampled. (**b**) Tracking the total size of CD8^+^ clones de-orphanized to each spike epitope within individual donors (thin grey lines). The mean of log_10_-transformed values across donors is also shown (thick blue lines). Red triangles indicate the timing of vaccination.

### Clonal succession in vaccine-induced CD8 T cell memory specific to B15-restricted spike epitopes

We previously reported that CD8^+^ T cell clones specific to the B15_NQK epitope, the most-studied spike epitope presented on HLA-B*15:01^31,36^, expand rapidly after the 1st vaccination but are not further boosted by the 2^nd^ or 3^rd^ doses^12^. Consistent with this, the B15_NQK-specific TCR cluster (cluster 16) is enriched for clones that peak after the 1^st^ dose (*1’ transient* or *1’ persistent*) (Fig. 6a,b). This behavior was somewhat unconventional and unexplained in previous reports^31,37^, so we focused on HLA-B*15:01^+^ individuals to characterize their overall response profiles. With the additional curation of VAS and FQP epitopes, we were able to identify robust B15 responses following a clear pattern of succession in epitope-specific responses.

**Fig. 6.**
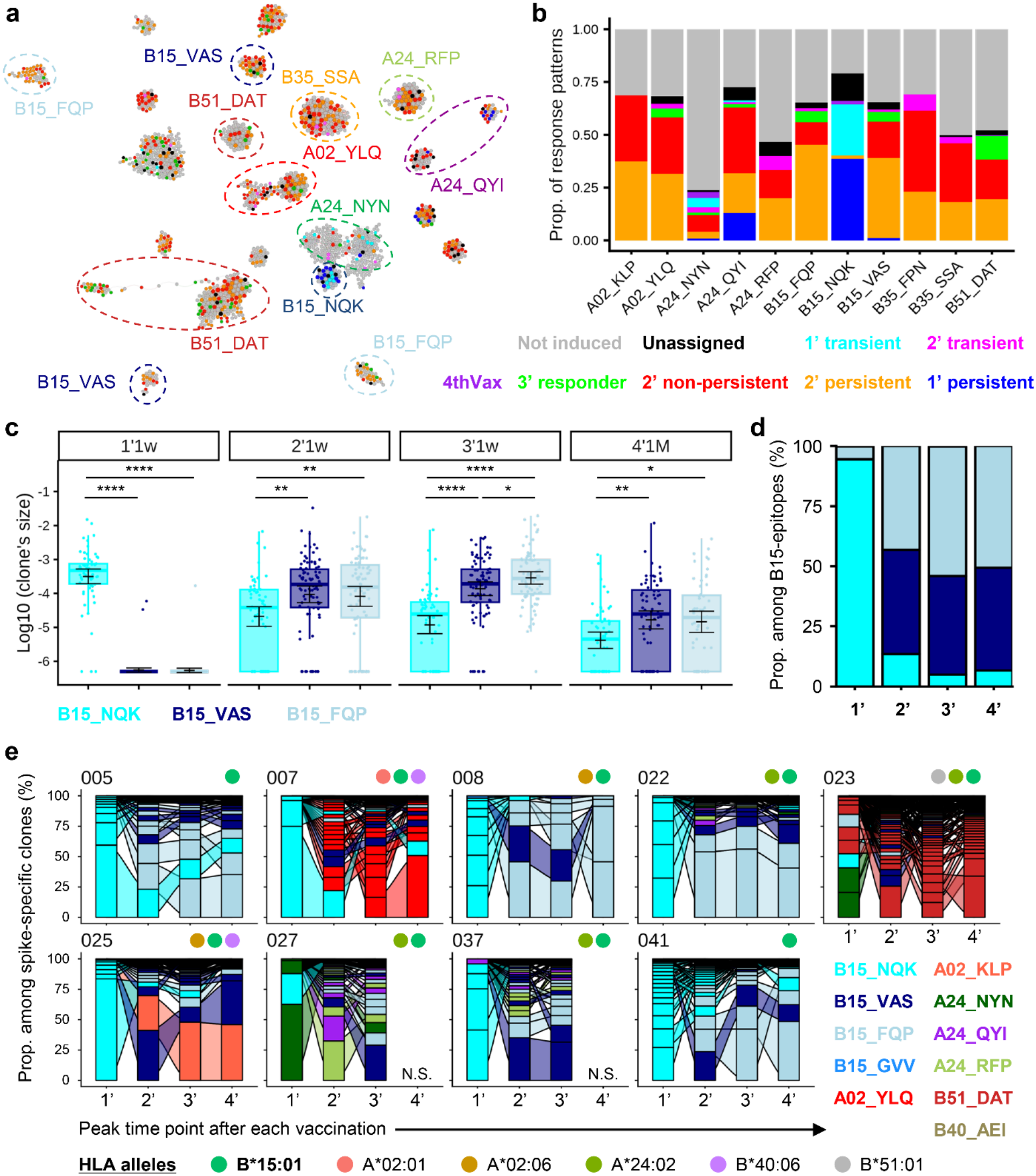
Longitudinal kinetics of CD8^+^ T cell clones specific to B15 spike epitopes. (**a**) TCR network graph of 25 major clusters colored with the response pattern of vaccine-associated clones, as defined in Fig. 1c. CD8^+^ clones that do not meet the criteria of vaccine-induced or that are not assigned to any response patterns are shown in grey or black, respectively. (**b**) Proportion of CD8^+^ clones showing each response pattern among those de-orphanized to each spike epitope. Only epitopes with more than ten de-orphanized clones are shown. *4thVax*; clones whose frequency increased more than 4-fold from baseline (P1) or exceeded 0.02% of the repertoire 1 week or 1 month after the 4^th^ dose. (**c**) Size of individual B15_NQK-, VAS- or FQP-specific clones after vaccinations in B*15:01^+^ donors (n=9). Horizontal lines and bars represent the mean and 95% CI of log_10_-transformed clones’ size, and box plots represent the median and the interquartile range. BH-adjusted p-values of Student’s t-test were shown. ****p<0.0001; ***p<0.001; **p<0.01; *p<0.05. NQK-specific clones; n=62. VAS-specific clones; n=87. FQP-specific clones; n=75. (**d**) Stacked bar plot showing the relative abundance of B15_NQK-, VAS-, and FQP-specific clones after vaccination, normalized to the total size of CD8 T cell clones recognizing B15-presented epitopes. Bar heights represent the mean proportions across HLA-B*15:01^+^ donors (n=9). (**e**) Sankey plots depicting trajectories of de-orphanized CD8^+^ vaccine-induced clones in donors carrying HLA-B*15:01. Each bar, connected by ribbons, represents an individual CD8^+^ T cell clone de-orphanized to a spike epitope. Bar height represents the proportion of each clone among those de-orphanized to spike epitopes, and bar colors correspond to respective epitopes. Numbers and colored circles above the plots denote individual donors and HLA alleles they carry. Only HLA alleles that present spike epitopes estimated for TCR clusters (summarized in Fig. 4d) are shown. 1’: 1 week after the 1^st^ dose. 2’: 1 week after the 2^nd^ dose. 3’: 1 week after the 3^rd^ dose. 4’: 1 month after the 4^th^ dose. N.S.: Not sampled.

Clusters specific to these other B15-spike epitopes (ex. clusters 13 and 24 for B15_VAS, and clusters 11 and 15 for B15_FQP) are enriched with *2’ persistent* and *2’ non-persistent* clones compared with the B15_NQK clusters (Fig. 6a,b). This led to a shift in the CD8^+^ clonal dominance among B15-presented spike epitopes (Fig. 6c,d): One week after the first dose B15_NQK-specific clones accounted for more than 90% of T cells targeting the B15-presented epitopes on average. However, this hierarchy reversed 1 week after the 2^nd^ dose, with FQP- and VAS-specific clones comprising more than 40%, and NQK-specific clones less than 15%, a pattern that persisted through 1 month after the 4^th^ dose (Fig. 6d). Consistently, within six of the nine B*15:01^+^ donors, the largest CD8^+^ T cell clones with an identified Spike target shifted from B15_NQK-specific clones (*cyan*) to B15_FQP- or VAS-specific clones (*light blue* or *navy*) after the 2^nd^ or 3^rd^ dose, visualized as crossing ribbons in the Sankey plots (Donors 005, 008, 022, 025, 037 and 041; Fig. 6e). In the remaining three donors, the largest to third-largest B15_NQK-specific clones after the 1^st^ dose ceded their position to clones targeting non-B15 epitopes after repeated vaccinations (Donor 007: to A02_YLQ-, Donor 023: to B51_DAT, Donor 027: to A24_RFP) and were also no longer the largest clones targeting B15-epitopes. Similar clonal successions were observed for other combinations of clones, such as from A24_NYN- or A24_QYI- to B51_DAT-specific clones in Donors 013, and 019 (Extended Data Fig. 7a-c).

Next, we examined possible mechanisms that could explain the clonal succession between NQK- and VAS/FQP-specific clones. First, NetMHCpan-4.1^33^ predicted that all three of these peptides are strong binders of HLA-B*15:01 (rank of predicted binding score: NQK: 0.12%, VAS: 0.116%, and FQP: 0.193%). This suggests that the observed clonal succession is unlikely to be explained by large differences in HLA binding affinity, which shifted our focus to the TCR-pHLA interaction itself.

To measure TCR-pHLA strength, we transduced NQK-, VAS-, and FQP-specific α/βTCRs into an NFAT-GFP reporter cell line and analyzed their reactivity using a co-culture assay. First, we compared the functional avidity of NQK-, VAS-, and FQP-specific TCRs by co-culturing them with HLA-B*15:01^+^ K562 cells in the presence of a dilution series of their target peptide. The resulting dose-response curves for GFP signals showed that FQP-and VAS-specific TCRs had higher functional avidity than NQK-specific TCRs (Fig. 7a,b). Since the sequence of the NQK epitope is well conserved between SARS-CoV-2 and other seasonal human coronaviruses (hCoVs) (Extended Data Fig. 8a), we also evaluated the cross-reactivity of NQK-specific TCRs to hCoV NQK peptides (Extended Data Fig. 8b). All five NQK TCRs examined recognized the NQK epitopes of SARS-CoV-2 and hCoVs (OC43 and HKU1) (Extended Data Fig. 8b,c).

**Fig. 7.**
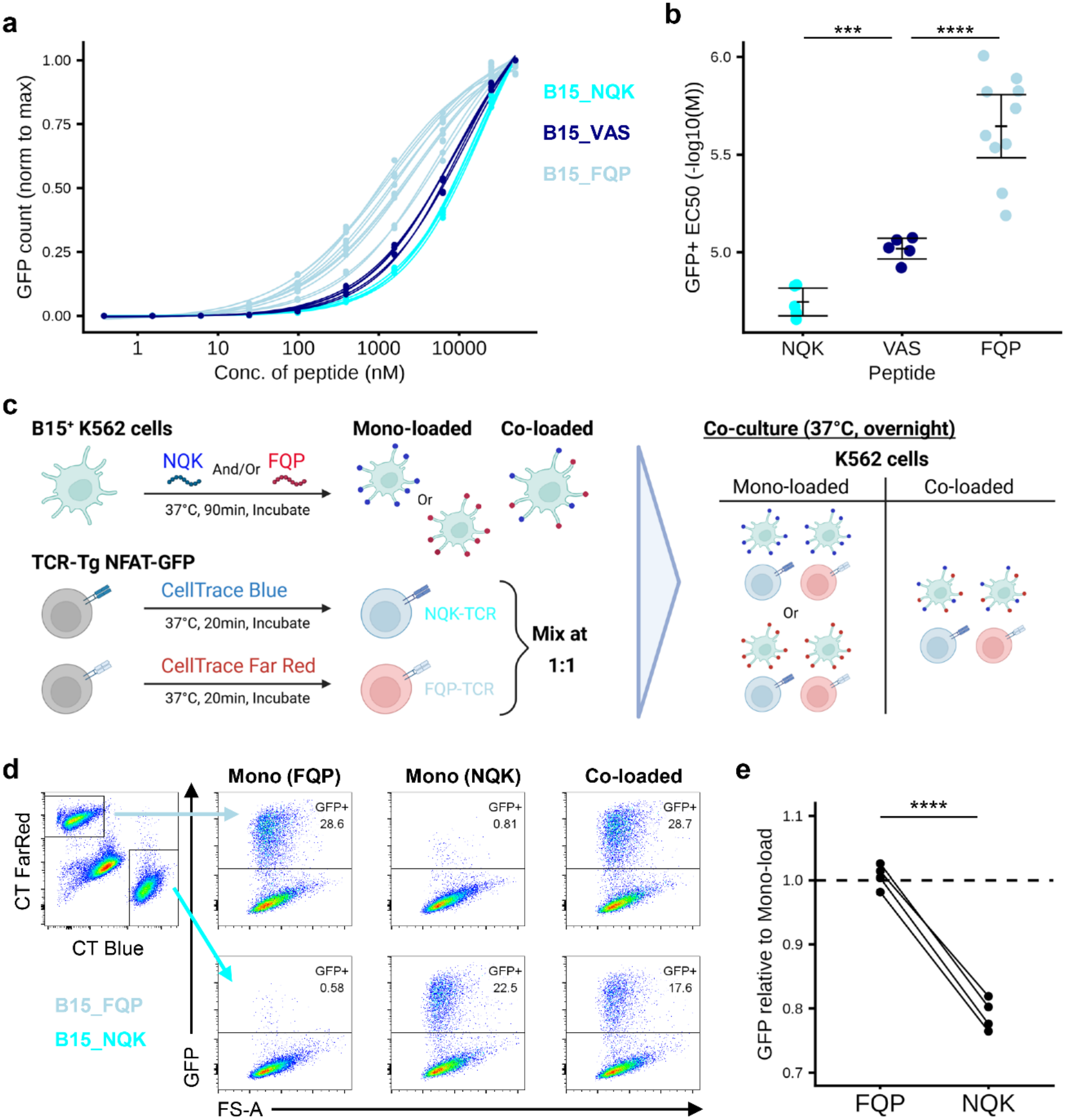
Avidity of vaccine-induced TCRs specific to B15 epitopes. (**a**) Dose-response curves for GFP signals from NFAT-GFP Jurkat cells transduced with NQK- (n=5), VAS- (n=5), or FQP-specific TCRs (n=10). Dots represent the GFP signals averaged between two independent experiments. (**b**) Comparing EC50 values between NQK-, VAS-, and FQP-specific TCRs. EC50 values are calculated as the peptide concentration corresponding to the half-maximal effective dose for each fitted dose–response curve. BH-adjusted p-values of the unpaired Student’s t-test were shown. (**c**) Schematic overview of the mixed co-culture experiment (*Method* in detail). (**d**) Gating strategy for evaluating GFP positivity in NQK- and FQP-specific TCR-transduced NFAT-GFP, which are labeled with CellTrace (CT) Blue or FarRed, respectively. The percentage of GFP^+^ cells in the FQP- or NQK-mono-loaded and co-loaded conditions is shown. (**e**) Competition effect between FQP- and NQK-specific TCRs. After subtracting the background GFP signal, the GFP positivity of NQK- or FQP-TCRs in co-loaded conditions was normalized to that of mono-loaded conditions and then compared between FQP- and NQK-TCRs for each combination (n=4: 2 NQK-TCRs x 2 FQP-TCRs). 6.25 μM of NQK and/or FQP peptide is loaded on B15^+^ K562 cells. P-values from the unpaired Student’s t-test are shown. ****p<0.0001; ***p<0.001; **p<0.01; *p<0.05. (**c**) Created using BioRender.com.

Interestingly, the strongest TCR activation was observed against SARS-CoV-2’s NQK peptide sequence (Extended Data Fig. 8c), indicating that the COVID-19 mRNA vaccine selectively mobilized cross-reactive CD8^+^ T cells that recognize the SARS-CoV-2 NQK epitope more strongly. These results suggest a hypothesis for the mechanism of clonal succession between B15_NQK- and B15_VAS/FQP-specific clones: B15_NQK-specific clones can rapidly respond to the first vaccination as pre-existing cross-reactive memory cells, with a substantially higher precursor frequency, while VAS- or FQP-specific clones are “de novo”, recruited from the naïve repertoire. However, their established memory precursor frequencies after two or more vaccinations and stronger epitope recognition allow for competition and domination of the NQK response in later exposures.

Next, we examined whether the FQP-specific TCRs outcompete NQK-specific TCRs in an in vitro setting. For this purpose, we mixed NQK- and FQP-specific TCR-transduced NFAT-GFP cells at a 1:1 ratio and co-cultured them with B15^+^ K562 cells loaded with an NQK or FQP peptide (*mono-loaded*) or both (*co-loaded*) (Fig. 7c). When co-cultured with NQK and FQP co-loaded K562 cells, both NQK and FQP TCR-transduced cells became GFP^+^ (Fig. 7d); however, the proportion of GFP^+^ cells among NQK TCR-transduced cells was reduced compared to the NQK-mono-loaded condition, while that of FQP TCR-transduced cells remained unchanged (Fig. 7e and Extended Data Fig. 8d). Moreover, by comparing single TCR vs. mono-loaded K562 (*single vs mono-loaded*) and single TCR vs. co-loaded K562 (*single vs co-loaded*), we were able to quantify the FQP vs. NQK TCR competition, with FQP TCRs reducing the GFP positivity of NQK TCRs relative to the *single vs mono-loaded* baseline (Extended Data Fig. 8e,f). These results indicate that high-avidity FQP-specific clones outcompete low-avidity NQK-specific clones for access to their antigens presented on the same APCs, providing a mechanistic explanation for clonal succession in B15-presented spike-specific CD8^+^ T cell memory during repeated vaccinations.

Finally, we examined whether mutations in these B15 spike epitopes observed in SARS-CoV-2 VOCs affected TCR recognition. No mutations are reported in the NQK and VAS epitopes for major VOCs. In contrast, the asparagine (N) at position 5 of the FQP epitope is mutated to tyrosine (Y) in VOCs, including Alpha, Beta, Gamma, and Omicron strains^38^, which has been associated with epitope escape^39^. Indeed, all 10 FQP-specific TCRs that we examined failed to recognize the mutated peptide (N5Y), even at high concentration (Extended Data Fig. 8g,h).

## Discussion

Although longitudinal TCR-seq studies have previously tracked the kinetics of T cell responses induced by SARS-CoV-2 mRNA vaccinations at clonotypic resolution^12,32,40^, the immunological interpretation of these data remained ambiguous because spike reactivities of vaccine-induced T cell clones were often not validated and, when assessed, were generally limited to the whole spike protein or a small number of epitopes. In this study, we performed TIRTL-seq to obtain repertoire-wide paired α/βTCR data from mRNA vaccinees and integrated it with longitudinal TCRβ-seq data from the same cohort. By combining several complementary methods for determining epitope specificity, including sequence similarity to known epitopes, structure-based modeling, and empirical reverse epitope discovery, followed by validation via TCR reconstitution in vitro, we expanded the epitope specificity landscape of vaccine-induced CD8^+^ T cell clones, representing 15 peptide epitopes presented on 8 HLA class I alleles, including 11 epitopes for which no cognate paired α/βTCR had previously been identified (Fig. 4d).

We found that, in donors carrying HLA-B*15:01, B15_NQK-specific T cell clones, which are cross-reactive to hCoVs, are successively replaced by clonotypes with higher-avidity TCRs recognizing VAS or FQP epitopes unique to SARS-CoV-2 after repetitive vaccinations. The stronger expansion of VAS-or FQP-specific clones in our dataset suggests that the T cell memory response in B*15:01^+^ donors should be monitored using VAS or FQP epitopes rather than the NQK epitope, especially after multiple exposures to SARS-CoV-2 spike. These clonal succession dynamics also indicate that epitope mapping based on early exposures may fail to identify immunodominant responses that emerge only after repeated boosting even with the same antigen. Notably, similar clonal succession dynamics were observed for other combinations of epitopes or clones, e.g. from A24_NYN to B51_DAT epitopes, which may also reflect a transition from cross-reactive to de novo T cell memory against SARS-CoV-2 spike. One remaining question is what types of spike-specific T cell memory are formed in donors who have not been exposed to coronaviruses and lack cross-reactive memory: are NQK-specific T cell responses suppressed by VAS-or FQP-specific T cells, or are high-avidity T cell clones specific to SARS-CoV-2 NQK induced? Analyzing vaccine-induced T cell responses in children born after the pandemic, who experience SARS-CoV-2 as their first coronavirus exposure, could answer this question and guide the timing of childhood SARS-CoV-2 vaccination to generate potent T cell memory targeting the conserved coronavirus epitope^41^.

After the finding that T cells recognizing the same epitope often have similar TCR sequences^22,23^, efforts have been devoted to predicting the epitopes of T cell clones from TCR sequence similarity to those with validated pHLA targets in public datasets^22,42^. More recently, 3D structures of TCR-pMHC complexes simulated by machine learning can also be used to predict T cell epitopes^43^. However, both sequence- and structure-based methods generally show limited accuracy when applied to previously unseen individual T cell clones without characterized reference epitopes. In this study, we first clustered vaccine-associated clones with similar TCRs, assuming they recognize the same epitope. Then, their target epitope was inferred at the cluster level by aggregating the predictions of individual clones using sequence or structure-based approaches within each cluster. This clustering-based approach leverages the coherence of TCR features within clusters to reduce noise from clone-level predictions.

Importantly, this approach provides epitope-specific resolution without the need to select epitopes to test before conducting experiments, as in pHLA multimer staining. The antigen-agnostic nature of the reverse epitope discovery TCR-clustering-based approach allowed us to analyze T cell responses to novel and less-characterized epitopes (e.g., B15_VAS and B15_FQP) and to integrate the latest updates in the database of spike epitopes and TCRs recognizing them (e.g., B51_DAT).

Despite these advances, the clustering-based approach could de-orphanize only 1,109 of 5,681 vaccine-associated clones at this point and depended on a rich auxiliary set of reference TCRs from SARS-CoV-2 post-exposure studies. One reason for this limitation is that about half of the vaccine-associated clones could not be clustered with others and remained as private clones. This raises the possibility that these stand-alone TCRs share structural similarity with “clustered” clones that is not captured by the TCR sequence similarity index used here (tcrdist). Notably, the specific spike epitopes of a subset of these stand-alone TCRs have already been identified by spike pHLA tetramer TCR-seq in our previous study^11,12^. Thus, comparing the structures of

TCR-pHLA ternary complexes between clustered and stand-alone clones may inform the development of improved TCR similarity metrics that better reflect their 3D structures. Another limitation is that clones recognizing epitopes presented on rare HLA alleles in our Japanese cohort are less likely to cluster. Incorporating paired α/βTCR data from more genetically diverse populations could expand the list of de-orphanized spike-reactive TCRs. Finally, we used conservative thresholds for clustering throughout our analysis to maximize specificity, which may have missed sequence convergence in some cases.

Despite these limitations, we provide a comprehensive dataset of α/βTCRs of CD8^+^ clones induced by SARS-CoV-2 mRNA vaccinations: these TCRs are linked to their longitudinal kinetics across repeated vaccinations and de-orphanized to one of the 15 spike epitopes. This resource enables understanding the broad landscape of mRNA vaccine-induced CD8^+^ responses at clonotypic resolution, linked to epitope specificities, and could support the development of improved algorithms for T cell epitope prediction based on TCR sequence similarities and structural modeling. The approach presented in this study paves the way for TCR-seq-based monitoring of antigen-specific T cell responses, offering a higher immunological resolution compared with T cell functional assays or pHLA multimer staining alone.

## Methods

### Study design

41 healthcare workers (≥20 years old) at Nara Medical University in Japan who had received the first dose of Pfizer-BNT162b2 vaccine in March 2021 were enrolled. Peripheral blood samples of 38 donors, collected at the following time points, were analyzed in this study: before receiving the first dose (P1), 1 week after the first dose (P2), 1 week (P3), 1 month (P4), 3 months (P5), and 6 months (P6) after the second dose, and 1 week (P8), 1 month (P9), 3 months (P10), and 6 months (P11) after the third dose. One participant (Donor 018) was not sampled at P11 because they had received the fourth vaccination before this time point. 23 of the participants received a fourth mRNA vaccination (either Pfizer–BNT162b2 or Moderna mRNA-1273) between August and December 2022, with samples obtained at ∼1 week, 1 month, 3 months, and 6 months post-vaccination (P13–P16), and additional annual samples in 2023 and 2024 (P17 and P18). PBMCs were isolated from whole blood with BD Vacutainer CPT tubes, resuspended in CELLBANKER 1 (Takara), and stored at -80°C. The detailed study design was described in Kitabatake et al.^10^ and Xu et al^11^.

### Study approval

The study was conducted in accordance with the Declaration of Helsinki and Good Clinical Practice Guidelines, following approval by the ethics board in each institution. All study participants provided written informed consent. The study was approved by the institutional review boards at Nara Medical University (Ethics Committee approval study No. 2954 and 3168), St. Jude Children’s Research Hospital (IRB study #24-1904), and Fred Hutchinson Cancer Center (IRB # FHIRB0021024).

### Classifying vaccine-induced clones into distinct response patterns using the longitudinal bulk TCRβ-seq dataset

Processed bulk TCRβ-seq dataset reported in our previous studies^11,12^, where each clonotype was defined by the CDR3 nucleotide sequence of their TCRβ, was re-analyzed to identify vaccine-induced clones. Vaccine-induced clones were initially defined as clonotypes whose frequency increased more than 4-fold from their baseline (P1) or exceeded 0.02% of the repertoire 1 week after the 1^st^, 2^nd^, or 3^rd^ vaccination. Then, these clones were classified into different response patterns using PCA and k-means clustering of their longitudinal kinetics. For PCA, each clone’s frequency at each time point was normalized to its maximum value across P1 to P11 and then log10-transformed after adding 1/1000 to accommodate missing values. Then, all calculated PC values of individual clones were used as inputs for k-means clustering. We chose k = 6 as the optimal number of clusters from the elbow plot. Vaccine-induced clones of the Donor 018 were excluded from the PCA analysis because this donor lacks the data at P11.

Vaccine-induced clones were reassigned to response patterns defined by PCA and k-means clustering. Representative kinetics for the six response patterns were calculated as the means of the normalized, log10-transformed frequency within each pattern. Then, Pearson correlation coefficients (R values) were calculated between each clone’s longitudinal kinetics and the representative kinetics of the six response patterns, and clones were re-assigned to the responding pattern with the highest R value. Clones with a maximum R < 0.7 were regarded as “unassigned” and excluded from vaccine-induced clones in the subsequent analyses. Vaccine-induced clones from Donor 018 (n=121) were assigned to response patterns in the same manner, except that R values were calculated without using the P11 time point.

### Building generalized linear models to classify *2’ persistent* and *2’ non-persistent* vaccine-induced clones based on their size.

Generalized linear models to classify the *2’ persistent* and *2’ non-persistent* vaccine-induced clones using their size at each time point were built. To determine when the longitudinal kinetics of these two groups of clones start being separated, we built multiple generalized linear models by adding the time points used from the beginning: the first model uses the time points before vaccination (P1) and 1 week after the 1^st^ dose (P2), the second model uses these two time points as well as the time point at 1 week after the 2^nd^ dose (P3), etc. Model performance was evaluated using the area under the receiver operating characteristic curve (ROC AUC) based on frequency-derived features.

### Re-analyzing single-cell data to obtain clonotype-level gene expression profiles

Single-cell TCR and targeted mRNA sequencing data obtained using the BD Rhapsody Immune Response Panel in our previous study^12^ were analyzed by Python scripts built on the *scanpy* package. The raw transcript counts for the 368 genes included in the panel were normalized to sum to 10,000, then log+1-transformed and averaged over all cells in each TCR clonotype. The clonotype-averaged, log-transformed, and normalized counts for these genes were standardized to have mean 0 and variance 1 with outlier values clipped at 10. After computing principal components (scanpy.tl.pca), batch effects across samples were corrected using Harmony (scanpy.external.pp.harmony_integrate). The top 15 corrected Harmony embeddings were then used for downstream UMAP dimensionality reduction (scanpy.tl.umap) and Leiden clustering (sc.pp.neighbors with n_neighbors=15, followed by scanpy.tl.leiden). Differentially expressed genes between *2’ persistent* and *2’ non-persistent* clones were identified using sc.tl.rank_genes_groups function with Wilcoxon test. Gene signature scores were calculated using sc.tl.score_genes function with marker genes for blood T cell subpopulations reported in Wang et al.^20^, and the Hallmark IFNα response gene set from Msigdb^21^. The list of genes used for calculating signature scores is summarized in Supplementary Table 6.

### Obtaining paired α/βTCRs with TIRTL-seq

CD4^+^ and CD8^+^ T cells were enriched from PBMC using Dynabeads CD4 and CD8 Positive Isolation Kit with bead detachment (11331D and 11333D Thermo Fisher), according to the manufacturer’s protocol. Enriched CD4^+^ and CD8^+^ cells are pooled for each sample and dispensed into 96 or 192 wells of 384-well plates with TIRTL-seq RT mixture, and TIRTL-seq libraries were prepared as described previously^14,44^. Final libraries were sequenced on Illumina NovaSeq platform at read length 2×150bp. Raw fastq files from each well were processed with mixcr v 4.6.0 using the *analyze* pipeline. Optional switches were set to *analyze:* the ‘genericamplicon’ preset with floating-left-alignment-boundary, floating-right-alignment-boundary for C segment and split-by-sample using tag pattern in Read1 (plate ID barcode). All other parameters were set to default. Tag-pattern switch was used to match and split the samples based on plate barcodes incorporated during the PCR I step. Processing steps included demultiplexing raw reads by plate barcode, aligning V, D and J segments, assembling identical reads into clonotypes and frequency-based error-correction.

TCR pairs were called from TIRTL-seq data as described previously using MAD-HYPE or T-SHELL algorithms with some modifications. First, to accommodate the smaller number of wells per sample, the T-SHELL threshold for calling TCR pairs was adjusted from the original paper to include the top 10th percentile of the second paired α chain that yields smaller adjusted p-values. Second, MAD-HYPE–called α–β pairs were excluded when the fraction of libraries in which both chains were detected was less than half of the libraries in which either the α or β chain was detected: (w_a_-w_ij_) < (w_a_+w_b_-w_ij_)*0.5 or (w_b_-w_ij_) < (w_a_+w_b_-w_ij_)*0.5, where *w_a_* and *w_b_* are the numbers of libraries in which the α and β chains were detected, respectively, and *w_ij_* is the number of libraries in which both were detected. Third, when a single TCRβ chain was paired with two or more TCRαs, we excluded the secondary α if (i) it had a lower MAD-HYPE score than the primary α, or (ii) its averaged relative frequency (“readFraction”) was less than 0.8 times that of the primary α.

### Integrating TIRTL-seq results with longitudinal bulk-seq

Paired αβ TCRs obtained by TIRTL-seq and single-cell TCR/RNA-seq performed in our previous study^12^ were merged into the donor-matched CD4^+^ and CD8^+^ longitudinal bulk TCRβ-seq datasets using the CDR3β nucleotide sequence as the clonotype identifier. When a CDR3β of paired α/βTCR was detected in both the CD4^+^ and CD8^+^ bulk TCRβ-seq data, likely due to the contamination of CD4 or CD8 T cells during enrichment from PBMC^12^, we assigned the clone to CD4 or CD8 according to which subset exhibited the higher peak frequency across time points. Then, TCRs that have the combination of TRAV1-2 and TRAJ33/AJ20/AJ12 or TRAV10 and TRAJ18 were excluded as Mucosal-Associated Invariant T cells (MAIT) or invariant Natural Killer T cells (iNKT), respectively^45,46^. This procedure yielded 105,880 CD8 and 287,095 CD4 paired αβTCRs with longitudinal kinetics following repetitive mRNA vaccinations in 38 donors from our cohort.

### Detection of statistically enriched groups of α/βTCRs with similar sequences (TCR clumps), which associated with mRNA vaccination

Detection of TCR clumps, statistically enriched groups of α/βTCR with sequence similarity, was performed as described in Schattgen et al.^27^. First, to identify significantly neighbor-enriched TCR sequences, background TCRα and TCRβ sequences were generated by shuffling V(D)J recombinations in the observed (“foreground”) sequences, then filtered to match key properties of the foreground sequences, such as CDR3 length, N-insertions, and V/J gene usages. For TCRα chains, V–J pairing distribution were also preserved to reflect known recombination biases.

Then, to estimate the likelihood of observing a given number of sequence neighbors for a TCR sequence, background TCRdist distributions were computed separately for α and β chains and then combined into a paired distribution, assuming random α/β pairing. This expected background TCRdist distance distribution was calculated for each observed TCR clonotype, and the probability of seeing an observed number of foreground neighbors with distance less than or equal to a given threshold T (24, 48, 72, and 96 tcrdist units) was estimated using the Poisson distribution. To account for multiple testing, the neighbor-count probabilities were multiplied by 4*C where C was the number of observed foreground clonotype sequences.

In the next step, these neighbor-enriched clonotypes were grouped into clusters of similar sequences, using a simple greedy graph-based clustering algorithm. A directed neighbor graph was constructed by looping over the four distance thresholds T in {24,48,72, and 96} and adding an edge from TCRi to TCRj if TCRi has a significantly-enriched neighbor number at threshold T and TCRdist(TCRi, TCRj) ≤ T. TCR sequence clusters were formed from this graph by repeatedly selecting the node with the highest number of outgoing edges as a new cluster center, assigning all of its connected neighbors as cluster members, and deleting the cluster center and all cluster members from the graph, preventing clusters from spreading too far across sequence space and increasing the likelihood of shared reactivity. Through these steps, 809 TCR clumps were identified from 105,880 paired α/βTCRs of CD8^+^ T cell clones.

In the last step, CD8^+^ TCR clumps associated with mRNA vaccination were defined as clumps containing at least three clones, including at least one vaccine-induced clone, and showing, on average in a clump, significant expansion after vaccination. Clump-level averaged clonal expansion was examined by testing whether frequencies within the clump at P2, P3, or P8 (1 week after 1^st^, 2^nd^, or 3^rd^ dose) exceeded a 4-fold increase over baseline (P1) using paired t-tests. 103 clumps whose minimum p-value across time points was less than 0.05 were defined as vaccine-associated clumps.

### Annotations of TCR epitope specificity using public database

Potential epitope specificity was assigned to individual TCRs as described in Schattgen et al^27^. A filtered version of the VDJdb^27–29^, added with SARS-CoV-2 spike-specific TCRs reported in previous studies^24–26,30–32^, was used as a reference database of TCRs with known pHLA target.

Matches were identified using the TCRdist measure, with statistical significance assigned by matching each TCR to a large, shuffled background database to create an expected probability distribution of TCRdist scores.

### Identifying mRNA-vaccine-associated α/βTCRs and clustering them

Vaccine-associated CD8^+^ α/βTCRs were defined as (I) TCRs identified as vaccine-induced in longitudinal bulk TCRβ-seq data (n = 4,473), or (II) TCRs belonging to vaccine-associated TCR clumps (n=2,142), yielding 5,681 unique TCRs. In donors monitored after the 4^th^ vaccination, we also considered clonotypes vaccine-induced if their frequency increased more than 4-fold from baseline (P1) or exceeded 0.02% of the repertoire 1 week or 1 month after the 4^th^ dose (“*4thVax*”). The sequence and longitudinal kinetics of 5,681 vaccine-associated TCRs are summarized in Supplementary Table 3. These 5,681 TCRs, together with 50 TCRs whose spike-specificity was validated, collected from 4 published studies^24–26,32^, were used as an input for *TCR clustering* analysis. First, pairwise TCRdist matrices were computed for all TCRs (n=5,731). Second, an undirected graph was generated using *igraph* R package^47^, with each node representing a unique TCR clone and edges connecting clone pairs with a TCRdist ≤ 72. Then, TCR clusters were identified using both connected component analysis and the Louvain community detection algorithm. This provided 25 major clusters containing 1,235 TCRs, 260 minor clusters containing 1,197 TCRs, and 3,299 singleton TCRs. A network graph of 25 major clusters was visualized using Gephi (v0.10.1)^48^. Amino acid sequence motifs (TCR logos) of paired αβTCRs for the 25 major clusters were created using *conga* python package^49^.

### Estimate targets of vaccine-associated TCR clusters via reverse epitope discovery

The HLA allele and/or spike epitopes were estimated for CD8^+^ TCR clusters containing more than 4 clones and observed in at least 2 donors using the following three independent methods.

#### Method 1: Using TCRs with validated spike-specificity

α/βTCRs with spike specificity validated in previous studies^12,24–26^ or annotated by the reference TCR database were counted for each clump. When a cluster contained at least three such validated TCRs, the corresponding pHLA target was estimated as the target of that cluster.

#### Method 2: Using HLA sharing across donors

For each CD8^+^ TCR cluster, we listed donors whose clones contributed to the cluster (“cluster’s donors”) and extracted their HLA-A and -B alleles reported in our previous study (“candidate HLA alleles”)^12^. Then, to test whether each candidate allele is significantly enriched among the cluster’s donors, we constructed a 2×2 contingency table comparing carriers vs. non-carriers among donors in the cluster to those in the rest of the cohort, and examined the enrichment using a one-sided hypergeometric test. The HLA allele with the smallest p-value among the candidates, provided it was less than 0.01, was designated as the predicted HLA allele associated with the cluster.

#### Method 3: Using public datasets of SARS-CoV-2 linked TCRβ

Two published datasets of TCRβs linked to SARS-CoV-2 exposure were used for this method. One is the dataset reported by Nolan et al.^35^, which contains TCRβs inferred to recognize SARS-CoV-2 spike epitopes identified by TCR-seq on CD8^+^ T cells activated by pooled peptide stimulation and subsequent antigen assignment (“*SARS-CoV-2 MIRA”*). The other is the Covid19-& HLA-associated TCR dataset^34^, which lists TCRβs that are more frequently observed in COVID-19-infected donors carrying a particular HLA allele. For each vaccine-associated TCR in our dataset, we identified TCRs in these two reference datasets that differed by at most one amino acid in the CDR3β sequence (“*edit-1 neighbors*”). Then, for each cluster, we determined the most frequent edit-1 neighbor. If this most frequent edit-1 neighbor is shared by more than 20% of clones in the cluster, its associated epitopes or HLA allele was estimated as the putative target of the cluster.

HLA alleles and epitopes independently predicted by the three methods were combined. For clusters with both an inferred HLA restriction and candidate target peptides, peptide–HLA binding affinities were predicted for them using NetMHCpan-4.1^33^, and only strong binders were retained as candidate epitopes. The prediction results for 25 major and remaining minor clusters were summarized in Supplementary Tables 4 and 5, respectively.

## Generate TCR-transduced cell lines

72 paired α/βTCRs from 13 clusters were selected for analyzing their reactivity to predicted spike epitopes (Supplementary Table 3). TCRα and TCRβ chains were modified to use murine constant regions (murine TRAC*01 and murine TRBC2*01). A gBlock gene fragment was synthesized by GenScript to encode the modified TCRα and TCRβ chains linked by 2A sites, which was cloned into the pLVX-EF1α-IRES-Puro lentiviral expression vector (Clontech). Then, these TCRs were transduced into TCR-null Jurkat 76.7 cells that express human CD8^+^ and an NFAT-GFP reporter construct (“*NFAT-GFP Jurkat cell line*”) using lentivirus vectors and subjected to antibiotic selection for 1 week using 1 µg ml^−1^ puromycin in RPMI (Gibco) containing 10% FBS and 1% penicillin–streptomycin, as reported in Minervina et al^24^.

Transduction was confirmed by evaluating surface TCR expression via flow cytometry using antibodies against mouse TCRβ constant region (1:100 dilution; Pacific Blue-conjugated, BioLegend 109226, clone H57-597) and human CD3 (1:20 dilution, APC-conjugated, BioLegend 344812, clone SK7).

### Peptide stimulation assay on TCR-transduced NFAT-GFP reporter cell line

To validate the specificity of vaccine-induced TCRs, 1×10^5^ of TCR-transduced NFAT-GFP Jurkat cells were co-cultured with 1×10^5^ of K562 cells (ATCC CCL-243) transduced with a single full-length HLA allele (HLA-A*02:01, HLA*A24:02, HLA-B*15:01, or HLA-B*51:01) or EBV-transformed lymphoblastoid cell lines (EBV-LCLs) carrying the matching HLA allele (HLA-B*35:01, or HLA-B*51:01; obtained from the Biospecimen Processing and Biorepository at the Fred Hutch Cancer Center) in a 96-well round-bottom plate, in the presence of the predicted target spike peptide purchased from GenScript (2 μM for A24_NYN-, A24_QYI-, and B35_SSA-specific TCRs, as well as B51_DAT-specific TCRs from the cluster 10, 50 μM for B51_DAT-specific TCRs from clusters 2 and 14, or 1.56 μM for the other TCRs). As a positive control, cells were stimulated with the eBioscience Cell Stimulation Cocktail at a 500-fold dilution. After overnight co-culture (20-24 hours), cells were resuspended, and one-third of the cell suspension was transferred to a 96-well flat-bottom plate. GFP signals from TCR-transduced NFAT-GFP Jurkat cells were quantified using an Incucyte live-cell analyzer with a 4-fold objective. Images were processed with Incucyte software (2025B and 2026A), and the count of GFP^+^ objects per image was used as the readout of TCR signaling.

### Quantifying the functional avidity of spike-specific TCRs using TCR-transduced NFAT-GFP reporter cell line

To evaluate the functional avidity of TCRs specific to spike epitopes presented on HLA-B*15:01 (NQK, VAS, and FQP), we co-cultured TCR-transduced NFAT-GFP Jurkat cells and B*15:01^+^ K562 cells, adding a dilution series of their corresponding peptides (50 μM, 25 μM, 25/4 μM, 25/4^2^ μM, … 25/4^6^ μM). The dose-response curves of TCR antigen recognition were generated by fitting four parameter logistic (4PL) models to GFP counts normalized to the maximum value within the peptide dilution series of each TCR. EC50 values are calculated as the peptide concentration that produces 50% of the maximal modeled GFP response for each fitted curve, using the “ED” function in the R package “*drc*”^50^, specifying a response level of 50%. Each TCR was evaluated in two independent experiments, each with two technical replicates; averaged values across technical replicates from both experiments were used to fit the logistic models.

### Structural modeling of TCR clusters and potential peptide epitopes

The *TCRdock* modeling pipeline^18^, updated to use the AlphaFold3 prediction network^17^, was used to build 3D models of TCR:pMHC complexes involving clustered TCRs. For each TCR cluster with a predicted HLA allele association and containing at least 10 TCRs, docking simulations were conducted against all 9mer spike epitopes that were predicted by NetMHCpan-4.1^33^ to be presented by the corresponding HLA allele. Since the number of clustered TCRs and predicted epitopes varied significantly across clusters, we performed a stochastic modeling procedure in which thousands of independent simulations were conducted, each docking a randomly selected cluster TCR against a randomly selected peptide epitope from the candidate list. The AlphaFold confidence in the predicted interface, as measured by the mean TCR:pMHC interchain ipTM score, was recorded for each simulation and potential peptides were ranked by the mean confidence over all independent docking simulations (typically 200-500 simulations per peptide). As an additional indicator of modeling confidence, the final TCR:pMHC docking geometries were recorded and compared across all models for each peptide epitope to assess the structural consistency of the ensemble of AlphaFold3 models (bar colors in Fig. 4b).

### Mixed co-culture assay with B15_NQK- and B15_FQP-specific TCRs

To evaluate the competition between B15_NQK- and B15_FQP-specific TCRs, we performed a mixed co-culture assay using these TCRs and B15^+^ K562 cells loaded with their target peptides. B15^+^ K562 cells were incubated for 90 minutes at 37 °C with either a NQK or an FQP peptide (*“mono-loaded”*) or both (*“co-loaded”*) to load the peptides onto their surface. The final concentration of each peptide was 25, 6.25, or 1.56 μM. NFAT-GFP reporter cells transduced with B15_NQK- and B15_FQP-specific TCRs were stained for 20 minutes at 37 °C in PBS containing CellTrace Blue (final 1 μM) or CellTrace FarRed (final 0.2 μM), respectively.

100,000 NQK- or FQP-specific NFAT-GFP cells (“*single TCR*”), or the same number of a 1:1 mixture of NQK- or FQP-specific NFAT-GFP cells (“*mixed TCRs*”), were co-cultured with 100,000 mono-loaded or co-loaded B15^+^ K562 cells. After an overnight co-culture, GFP positivity within the CellTrace Blue or FarRed^+^ fraction was measured by flow cytometry. After subtracting background GFP signals from the unloaded condition, the competition effect between NQK- and FQP-TCRs was calculated using two methods. In one way, the competition effect was calculated as the relative GFP positivity in the co-loaded condition compared to the mono-loaded condition (Fig. 7e and Extended Data Fig. 8d). In another way, the reduction in GFP positivity in *single vs co-loaded*, *mixed vs mono-loaded*, and *mixed vs co-loaded* conditions from *single vs mono-loaded* was calculated, and then the effect of inter-TCR competition was evaluated as (% reduction in *mixed vs co-loaded*) - (% reduction in *mixed vs mono-loaded*) - (% reduction in *single vs co-loaded*) (Extended Data Fig. 8f).

## Statistical analysis

Statistical significance was calculated using R v.4.4.0. Statistical tests used are stated in the respective figure legends. *P* values are labeled in each figure. TCR network graphs in Fig. 3b-e and Fig. 6a were created with Gephi (ver 0.10). Alignment of spike B15_NQK epitope across SARS-CoV-2 and other hCoVs was performed using SnapGene (ver 8.2.1).

## Supplementary Materials

Extended Data Fig. 1. Unsupervised analysis of longitudinal kinetics of vaccine-induced CD8 clones.

Extended Data Fig. 2. Comparison between *2’ persistent* and *2’ non-persistent* vaccine-induced CD8^+^ clones.

Extended Data Fig. 3. Rescuing CD8^+^ vaccine-associated clones with weaker clonal expansion

Extended Data Fig. 4. Structural modeling of TCR clusters and potential peptide epitopes.

Extended Data Fig. 5. Experimental validation of spike epitopes of TCR clusters.

Extended Data Fig. 6. Total size of T cell clones with or without predicted spike epitopes after each vaccination dose.

Extended Data Fig. 7. Longitudinal kinetics of vaccine-associated clones de-orphanized to spike epitopes in B*15:01^-^ donors.

Extended Data Fig. 8. Characterization of SARS-CoV-2 spike-specific TCRs targeting B*15:01-presented epitopes by in vitro co-culture assay.

Supplementary Table 1. Metadata of donors and sampling

Supplementary Table 2. List of CD8^+^ vaccine-induced clones identified in longitudinal bulk TCRβ-sequencing dataset.

Supplementary Table 3. List of paired α/βTCRs associated with mRNA vaccination. Their predicted/validated target spike epitopes, associated HLA alleles, and longitudinal kinetics are also summarized.

Supplementary Table 4. Predicted peptide and HLAs for 25 major TCR clusters of CD8^+^ vaccine-associated clones.

Supplementary Table 5. Predicted peptide and HLAs for minor TCR clusters of CD8^+^ vaccine-associated clones.

Supplementary Table 6. List of genes used for calculating signature scores in Fig. 2k.

## Acknowledgements

We thank the Hartwell Center Team at St. Jude and, in particular, S. Olsen, G. Neale and D. Darnell for their help with high-throughput sequencing; the Flow Cytometry Shared Resource, (RRID: SCR022613), the Cellular Imaging Shared Resource (RRID: SCR022609) of the Fred Hutch/University of Washington/Seattle Children’s Cancer Consortium (NIH P30 CA015704), and H. Abe and ImmunoGeneTeqs, Inc., for their help with the bulk TCR sequencing. Fig. 1a, 3a, 4a, 7c, Extended Data Fig. 1a, and 8e were created with BioRender. Perplexity AI and Grammarly were used in preparing the manuscript. The authors reviewed and verified all AI-assisted text, citations, and references, and take full responsibility for the final content of the manuscript.

## Author contributions

Conceptualization: H.A., M.V.P., A.A.M., S.U., K.M., and P.G.T.

Methodology: H.A. and M.V.P.

Investigation: H.A., Z.B.S., and V.B.

Formal analysis: H.A., M.V.P., P.B., A.A.M., and K.V.

Software: H.A., M.V.P., P.B., and K.V.

Data curation: H.A., E.K.A., M.K., N.O.S., and T.I.

Visualization: H.A. and P.B.

Resources: C.A., K.M.B., S.A.S., P.X., C.M., F.S., A.L., S.U., and T.I.

Supervision: P.G.T.

Funding acquisition: C.M., S.U., K.M., and P.G.T.

Writing, original draft: H.A.

Writing, review, and editing: all authors.

## Data availability

Raw sequencing data of TIRTL-seq are available via the Sequence Read Archive (SRA) (PRJNA1529769).

## Code availability

Code and data used in this study are available via Zenodo at https://doi.org/10.5281/zenodo.21881944.

## References

1. Altmann, D. M. & Boyton, R. J. COVID-19 vaccination: the road ahead. Science 375, 1127–1132 (2022).

2. Han, X., Xu, P. & Ye, Q. Analysis of COVID-19 vaccines: types, thoughts, and application. J. Clin. Lab. Anal. 35, e23937 (2021).

3. Kedzierska, K. & Thomas, P. G. Count on us: T cells in SARS-CoV-2 infection and vaccination. Cell Rep. Med. 3, 100562 (2022).

4. Moss, P. The T cell immune response against SARS-CoV-2. Nat. Immunol. 23, 186–193 (2022).

5. Gao, Y. et al. Ancestral SARS-CoV-2-specific T cells cross-recognize the Omicron variant. Nat. Med. 28, 472–476 (2022).

6. Painter, M. M. et al. Rapid induction of antigen-specific CD4+ T cells is associated with coordinated humoral and cellular immunity to SARS-CoV-2 mRNA vaccination. Immunity 54, 2133–2142.e3 (2021).

7. Guerrera, G., et al. BNT162b2 vaccination induces durable SARS-CoV-2-specific T cells with a stem cell memory phenotype. Sci. Immunol. 6, eabl5344 (2021).

8. Saini, S. K., et al. SARS-CoV-2 genome-wide T cell epitope mapping reveals immunodominance and substantial CD8+ T cell activation in COVID-19 patients. Sci. Immunol. 6, eabf7550 (2021).

9. Amaya Hernandez, S. P. et al. Comprehensive longitudinal profiling of SARS-CoV-2-specific CD8+ T-cells reveal strong functional impairment and recognition bias as markers for disease severity. Preprint at 10.1101/2025.03.25.644515 (2025).

10. Kitabatake, M. et al. Transition of antibody titers after SARS-CoV-2 mRNA vaccination in Japanese healthcare workers. Jpn. J. Infect. Dis. 76, 72–76 (2023).

11. Xu, P. et al. Repeated mRNA vaccination and breakthrough infection reveal durable dominance but diminishing recall of vaccine-induced CD8+ T-cell clones. Int. Immunol. 38, 371–382 (2026).

12. Aoki, H. et al. CD8+ T cell memory induced by successive SARS-CoV-2 mRNA vaccinations is characterized by shifts in clonal dominance. Cell Rep. 43, 113887 (2024).

13. Roland, M. M. & Joglekar, A. V. TIRTL-seq: heroes with T-SHELL. Nat. Methods 23, 16–17 (2026).

14. Pogorelyy, M. V. et al. TIRTL-seq: deep, quantitative and affordable paired TCR repertoire sequencing. Nat. Methods 23, 56–64 (2026).

15. Turchaninova, M. A. et al. Pairing of T-cell receptor chains via emulsion PCR. Eur. J. Immunol. 43, 2507–2515 (2013).

16. Seitz, S., et al. Reconstitution of paired T cell receptor α- and β-chains from microdissected single cells of human inflammatory tissues. Proc. Natl Acad. Sci. USA 103, 12057–12062 (2006).

17. Abramson, J. et al. Accurate structure prediction of biomolecular interactions with AlphaFold 3. Nature 630, 493–500 (2024).

18. Bradley, P. Structure-based prediction of T cell receptor:peptide-MHC interactions. eLife 12, e82813 (2023).

19. Blaser, C. et al. β-galactoside-binding protein secreted by activated T cells inhibits antigen-induced proliferation of T cells. Eur. J. Immunol. 28, 2311–2319 (1998).

20. Wang, Y. et al. Integrating single-cell RNA and T cell/B cell receptor sequencing with mass cytometry reveals dynamic trajectories of human peripheral immune cells from birth to old age. Nat. Immunol. 26, 308–322 (2025).

21. Liberzon, A. et al. The Molecular Signatures Database hallmark gene set collection. Cell Syst. 1, 417–425 (2015).

22. Glanville, J. et al. Identifying specificity groups in the T cell receptor repertoire. Nature 547, 94–98 (2017).

23. Dash, P. et al. Quantifiable predictive features define epitope-specific T cell receptor repertoires. Nature 547, 89–93 (2017).

24. Minervina, A. A. et al. SARS-CoV-2 antigen exposure history shapes phenotypes and specificity of memory CD8+ T cells. Nat. Immunol. 23, 781–790 (2022).

25. Nakama, T. et al. Position-5-driven reorientation of an immunodominant HLA-A24:02 SARS-CoV-2 epitope drives universal T cell escape. JCI Insight 11, e202235 (2026).

26. Kocher, K., et al. Vaccination-induced T cell responses maintain polyclonality with high antigen receptor avidity. Sci. Immunol. 10, eadu6730 (2025).

27. Schattgen, S. et al. Diverse modes of T cell receptor sequence convergence define unique functional and cellular phenotypes. Preprint at 10.1101/2025.05.31.657155 (2025).

28. Goncharov, M. et al. VDJdb in the pandemic era: a compendium of T cell receptors specific for SARS-CoV-2. Nat. Methods 19, 1017–1019 (2022).

29. Shugay, M. et al. VDJdb: a curated database of T-cell receptor sequences with known antigen specificity. Nucleic Acids Res. 46, D419–D427 (2018).

30. Pogorelyy, M. V. et al. Resolving SARS-CoV-2 CD4+ T cell specificity via reverse epitope discovery. Cell Rep. Med. 3, 100697 (2022).

31. Rowntree, L. C., et al. HLA-B15:01-positive severe COVID-19 patients lack CD8+ T cell pools with highly expanded public clonotypes. Proc. Natl Acad. Sci. USA 122, e2503145122 (2025).

32. Ford, E. S. et al. Repeated mRNA vaccination sequentially boosts SARS-CoV-2-specific CD8+ T cells in persons with previous COVID-19. Nat. Immunol. 25, 166–177 (2024).

33. Alvarez, B. et al. NNAlign-MA; MHC peptidome deconvolution for accurate MHC binding motif characterization and improved T-cell epitope predictions. Mol. Cell. Proteomics 18, 2459–2477 (2019).

34. Mayer-Blackwell, K. et al. TCR2HLA: calibrated inference of HLA genotypes from TCR repertoires enables identification of immunologically relevant metaclonotypes. PLoS Comput. Biol. 22, e1013767 (2026).

35. Nolan, S. et al. A large-scale database of T-cell receptor beta sequences and binding associations from natural and synthetic exposure to SARS-CoV-2. Front. Immunol. 16, 1488851 (2025).

36. Augusto, D. G. et al. A common allele of HLA is associated with asymptomatic SARS-CoV-2 infection. Nature 620, 128–136 (2023).

37. Oberhardt, V. et al. Rapid and stable mobilization of CD8+ T cells by SARS-CoV-2 mRNA vaccine. Nature 597, 268–273 (2021).

38. Scarpa, F. & Ciccozzi, M. Genetic variability of the SARS-CoV-2 JN.1 lineage. Pathog. Glob. Health 118, 277–279 (2024).

39. Le Bert, N. et al. Mutations of SARS-CoV-2 variants of concern escaping Spike-specific T cells. Preprint at 10.1101/2022.01.20.477163 (2022).

40. Ren, C. et al. Longitudinal change trend of the TCR repertoire reveals the immune response intensity of the inactivated COVID-19 vaccine. Mol. Immunol. 163, 39–47 (2023).

41. Benjamin Hills, L., et al. Pediatric T cell and B cell responses to SARS-CoV-2 infection. JCI Insight 10, e196032 (2025).

42. Mayer-Blackwell, K. et al. TCR meta-clonotypes for biomarker discovery with tcrdist3 enabled identification of public, HLA-restricted clusters of SARS-CoV-2 TCRs. eLife 10, e68605 (2021).

43. Dratva, L. M., Teichmann, S. A. & Kretschmer, L. Decoding human T cell immunity with artificial intelligence and single-cell genomics. Annu. Rev. Immunol. 44, 381–405 (2026).

44. Baretti, M. et al. A therapeutic peptide vaccine for fibrolamellar hepatocellular carcinoma: a phase 1 trial. Nat. Med. 31, 4246–4255 (2025).

45. Held, K., Beltrán, E., Moser, M., Hohlfeld, R. & Dornmair, K. T-cell receptor repertoire of human peripheral CD161hiTRAV1-2+ MAIT cells revealed by next generation sequencing and single cell analysis. Hum. Immunol. 76, 607–614 (2015).

46. Greenaway, H. Y. et al. NKT and MAIT invariant TCRα sequences can be produced efficiently by VJ gene recombination. Immunobiology 218, 213–224 (2013).

47. Csárdi, G. & Nepusz, T. The igraph software package for complex network research. InterJournal Complex Systems, 1695 (2006).

48. Bastian, M., Heymann, S. & Jacomy, M. Gephi: an open source software for exploring and manipulating networks. Proc. Int. AAAI Conf. Web Soc. Media 3, 361–362 (2009).

49. Schattgen, S. A. et al. Integrating T cell receptor sequences and transcriptional profiles by clonotype neighbor graph analysis (CoNGA). Nat. Biotechnol. 40, 54–63 (2022).

50. Ritz, C., Baty, F., Streibig, J. C. & Gerhard, D. Dose-response analysis using R. PLoS ONE 10, e0146021 (2015).

