## Supplementary Figures and Tables for "Repeated exposure to a conserved antigen drives dynamic clonal and epitope succession"

Extended Data Fig. 1

**a**

**Identify vax-induced clones** as

1. > 4-fold increase from the baseline (P1).
2. > 0.02%.
3. At 1w after 1st, 2nd, or 3rd vaccination.

**Define response patterns of clones** based on kinetics

1. Normalize clone's size at each point to its maximum.
2. Transform to log10 scale.
3. Perform PCA and k-means clustering.

**Re-assign clones to response patterns** by

1. Calculate R value with representative PCA kinetics.
2. Re-assign to the pattern with highest R value
3. Filter out clones if their R value < 0.7 ("unassigned")

**Compare response patterns** up to 2 years after 4th vax. in terms of

1. their total size within a donor
2. proportion of maintained clones (= breadth)

**b**

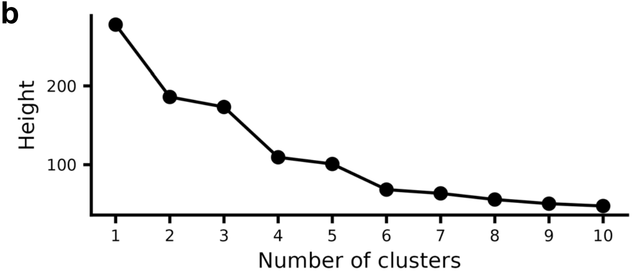


**c**

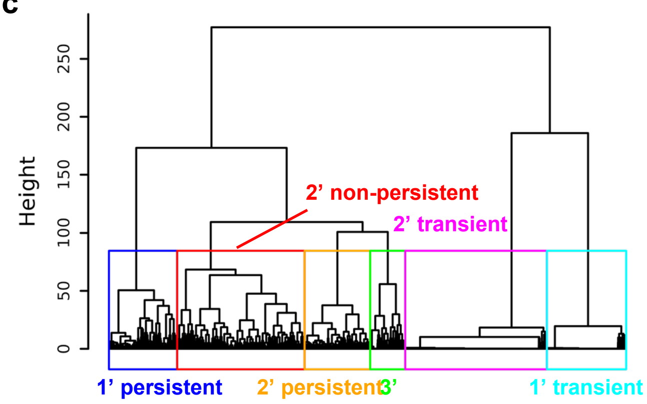


**d** **e** **f**

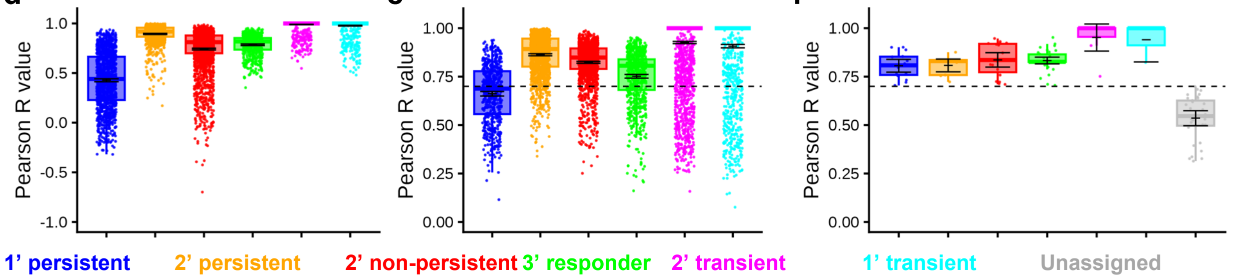


**g**

**After re-assignment**

Assigned by PCA

| | 1' persistent | 2' persistent | 2' non-persistent | 3' | 2' transient | 1' transient | Unassigned |
|---|---|---|---|---|---|---|---|
| 1' persistent | 261 | 32 | 34 | 28 | 29 | 12 | 895 |
| 2' persistent | 0 | 1161 | 18 | 15 | 1 | 0 | 46 |
| 2' non-persistent | 19 | 244 | 1533 | 8 | 182 | 48 | 377 |
| 3' | 0 | 15 | 0 | 529 | 0 | 0 | 120 |
| 2' transient | 0 | 0 | 1 | 0 | 2669 | 0 | 11 |
| 1' transient | 0 | 0 | 0 | 0 | 0 | 1469 | 31 |
| **Total** | **280** | **1452** | **1586** | **580** | **2881** | **1529** | **1480** |

**h**

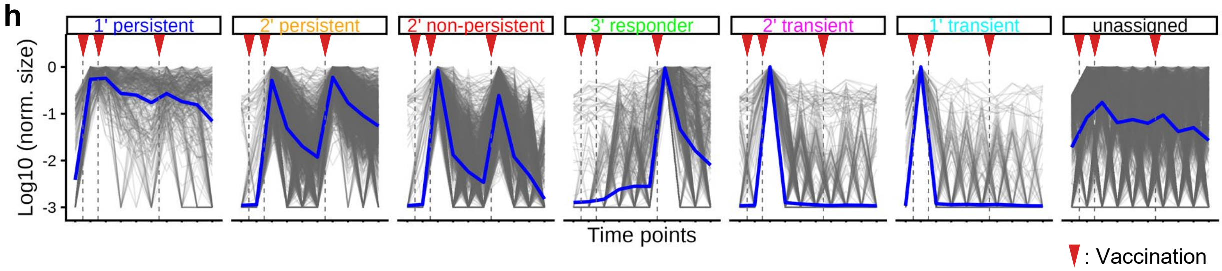

**Extended Data Fig. 1. Unsupervised analysis of longitudinal kinetics of vaccine-induced CD8 clones.**

(**a**) Workflow for classifying $CD8^+$ vaccine-induced clones into distinct response patterns. (**b**) Elbow plot for k-means clustering on vaccine-induced clones based on their longitudinal kinetics. (**c**) Dendrogram of longitudinal kinetics of vaccine-induced clones. (**d,e**) Pearson correlation coefficients (R values) between clones' longitudinal kinetics and the representative kinetics of the response patterns. The R values with the originally assigned response pattern (**d**) and those with the best-matching pattern (highest R among all six patterns) (**e**) were shown. (**f**) R values between the longitudinal kinetics of vaccine-induced clones and the best-matching pattern among the six patterns in Donor 018, who lacks the sample at P11. Clones with maximum $R < 0.7$ were regarded as "unassigned" and excluded from vaccine-induced clones in the subsequent analyses. (**g**) Contingency table showing the number of vaccine-induced clones originally assigned to six response patterns (rows) vs. after the reassignment based on their highest R values (columns). Vaccine-induced clones from Donor 018 were excluded from this table. (**h**) Normalized and $\log_{10}$-transformed kinetics of individual vaccine-induced clones classified into one of the six response patterns or left unassigned. Blue lines represent the mean values for each response pattern. (**a**) Created using BioRender.com.

Extended Data Fig. 2

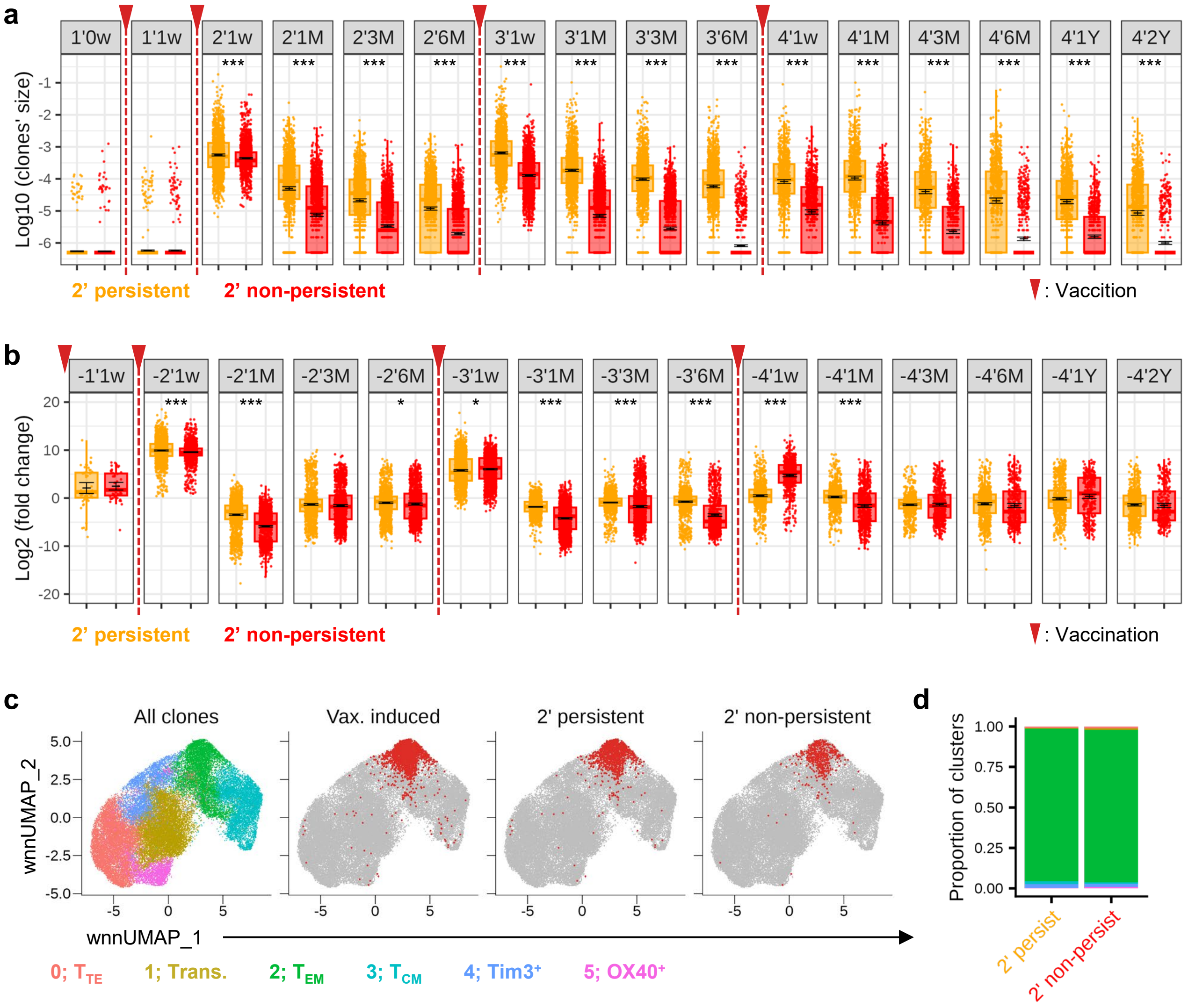


**Extended Data Fig. 2. Comparison between *2’ persistent* and *2’ non-persistent* vaccine-induced CD8$^+$ clones.**
(**a**) Size of individual *2’ persistent* and *2’ non-persistent* clones at each time point. (**b**) Fold changes in the size of individual *2’ persistent* and *2’ non-persistent* clones between each interval. The mean and 95% CI across clones, as well as the fold difference in the mean between the two groups, are shown. *2’ persistent*; n=1,452 clones. *2’ non-persistent*; n=1,586 clones. (**c**) Cell-level UMAP of gene expression. Each dot represents a CD8$^+$ T cell. Cells belonging to vaccine-induced clones (left), as well as *2’ persistent* (middle) and *2’ non-persistent* (right) clones at 1 week after the 2$^{nd}$ dose, are also shown. (**d**) Bar plot representing the distribution of CD8$^+$ T cells belonging to *2’ persistent* and *2’ non-persistent* clones at 1 week after the 2$^{nd}$ dose. P values of Student’s t-test were shown. ***p<0.001; *p<0.05. (**a,b**) Clones from Donor 018, who lacks data at P11, were excluded.

Extended Data Fig. 3

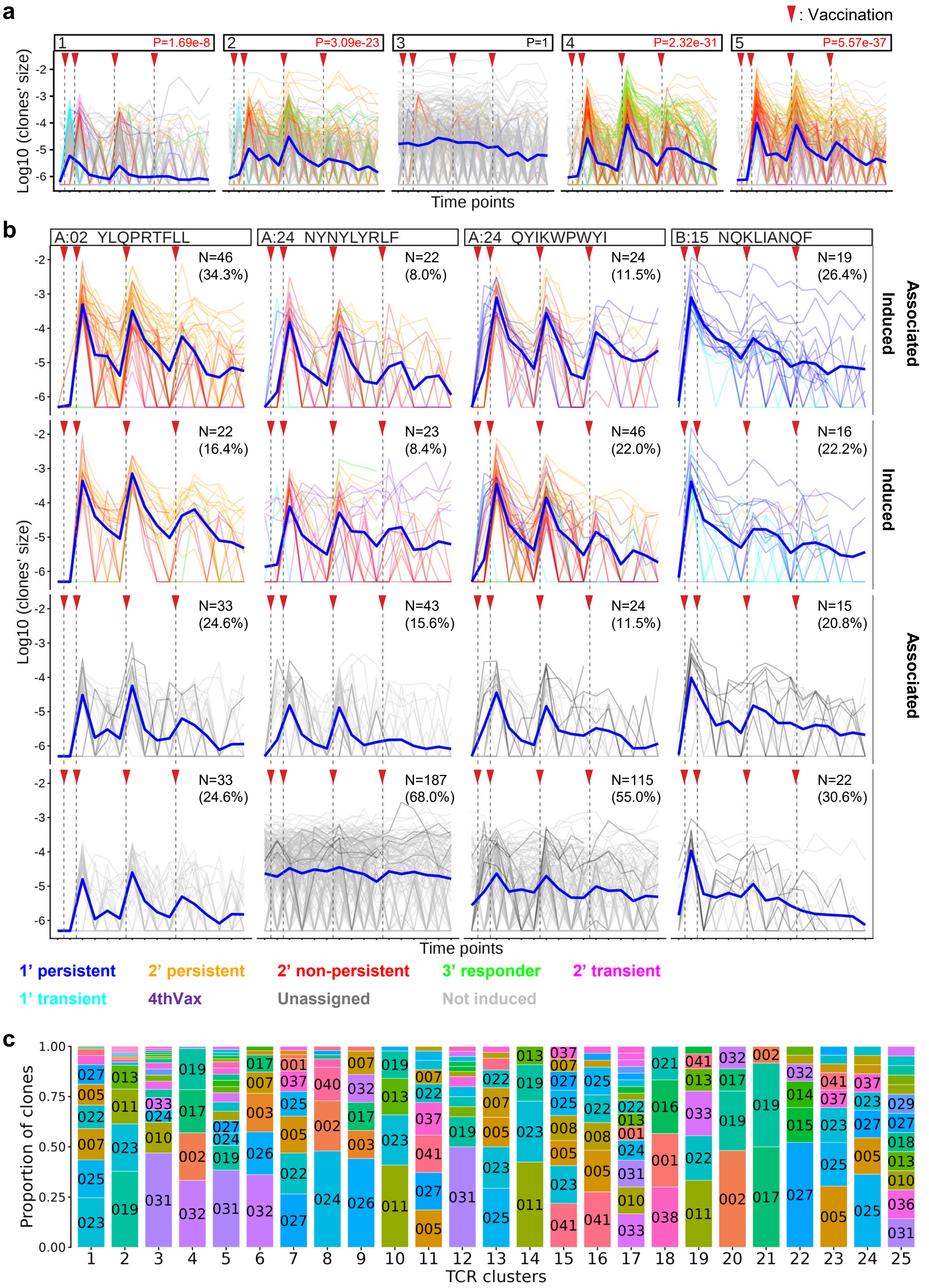

**Extended Data Fig. 3. Rescuing CD8$^+$ vaccine-associated clones with weaker clonal expansion.**

(**a**) Kinetics of CD8$^+$ clones included in the five representative TCR clumps classified as vaccine-associated or non-associated. A paired t-test was performed to assess whether clones within each clump expanded more than 4-fold from baseline to post-vaccination time point (after the 1$^{st}$, 2$^{nd}$, or 3$^{rd}$ dose), and the smallest p-value across doses is shown for each clump. In these examples, clumps 1, 2, 4, and 5 are classified as vaccine-associated, while clump 3 is not. (**b**) Kinetics of CD8$^+$ clones identified by spike pHLA tetramer TCR-seq for four epitopes (A02_YLQ, A24_NYN, A24_QYI, and B15_NQK), as described previously (*11*, *12*). pHLA tetramer$^+$ clones were classified into four groups based on whether their longitudinal kinetics fulfill the definition of vaccine-induced clones ("induced") or whether they were included in TCR clumps associated with vaccination ("associated"). The number and proportion of clones within each epitope specificity are shown. Lines represent the individual CD8$^+$ clones, colored by their response patterns, as defined in Fig. 1c. *4thVax*: clones whose frequency increased more than 4-fold from baseline (P1) or exceeded 0.02% of the repertoire 1 week or 1 month after the 4$^{th}$ dose. (**c**) Donor composition of clones within each major TCR cluster. Each color represents an individual donor, and donor IDs are shown for blocks comprising more than 6% of clones within a cluster.

### Extended Data Fig. 4

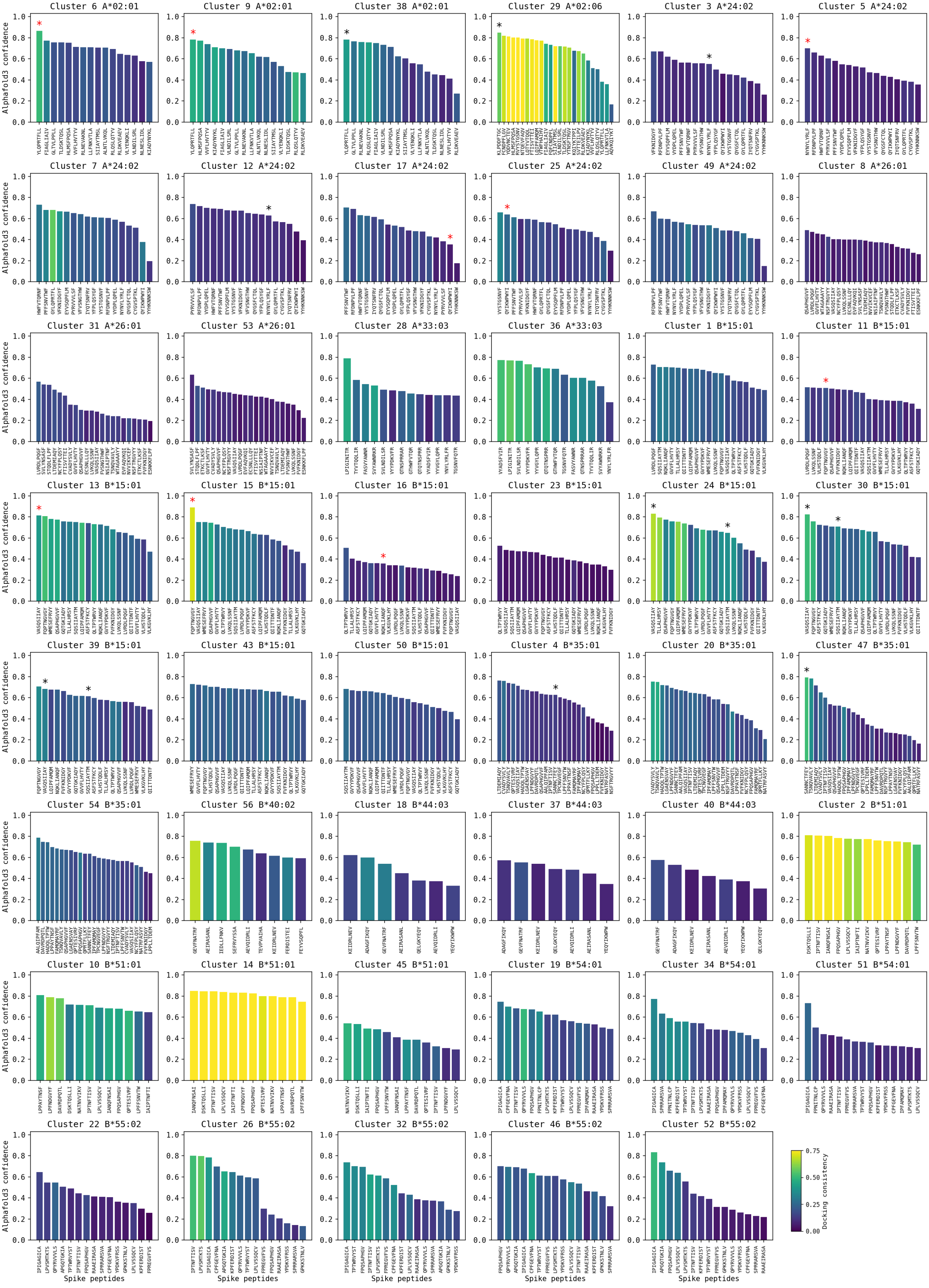

**Extended Data Fig. 4. Structural modeling of TCR clusters and potential peptide epitopes.**
Structure-based peptide prediction for TCR clusters whose associated HLA alleles are identified from donors' HLA typing results. 3D models of TCR:pMHC complexes for 9-mer spike epitopes predicted to be strong binders to associated HLA alleles were generated using the *TCRdock* pipeline with the AlphaFold3 prediction network. AlphaFold confidence in the predicted interface, measured by the mean TCR:pMHC interchain ipTM score, is shown as bars. Bar colors indicate the consistency of modeled TCR:pMHC docking geometries across TCRs, an additional indicator of modeling confidence. Asterisks show the estimated (in black) or experimentally confirmed (in red) target spike epitopes of the clusters (Supplementary Tables 4 and 5).

Extended Data Fig. 5

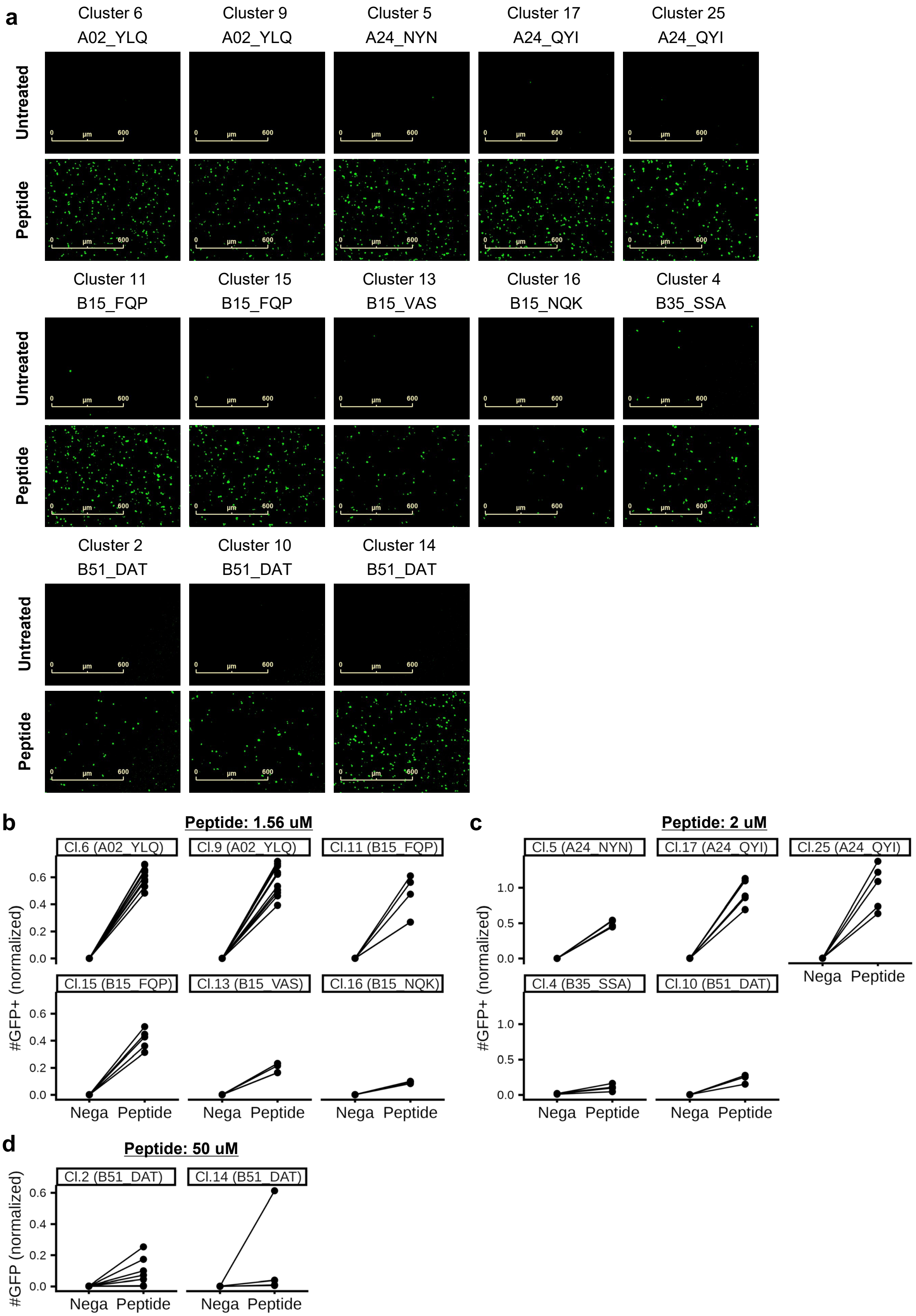

**Extended Data Fig. 5. Experimental validation of spike epitopes of TCR clusters.** (**a**) Representative Incucyte images showing GFP signals from TCR-transduced NFAT-GFP reporter cells after the co-culture assay. TCR-transduced NFAT-GFP cells were co-cultured with EBV-LCLs carrying the matching HLA allele (for B35_SSA-specific TCRs and B51_DAT-specific TCRs from clusters 2 and 14) or K562 cells transduced with the matching HLA allele (for other TCRs), with or without their target spike peptides. (**b-d**) The count of $GFP^+$ objects per image, normalized to the positive control stimulated with the Cell Stimulation cocktail. Each pair of dots connected by a line represents an individual TCR tested under untreated ("Nega") and peptide-stimulated conditions ("Peptide"). Target peptides were added at 1.56 μM (**b**), 2 μM (**c**), or 50 μM (**d**) to the co-culture.

Extended Data Fig. 6

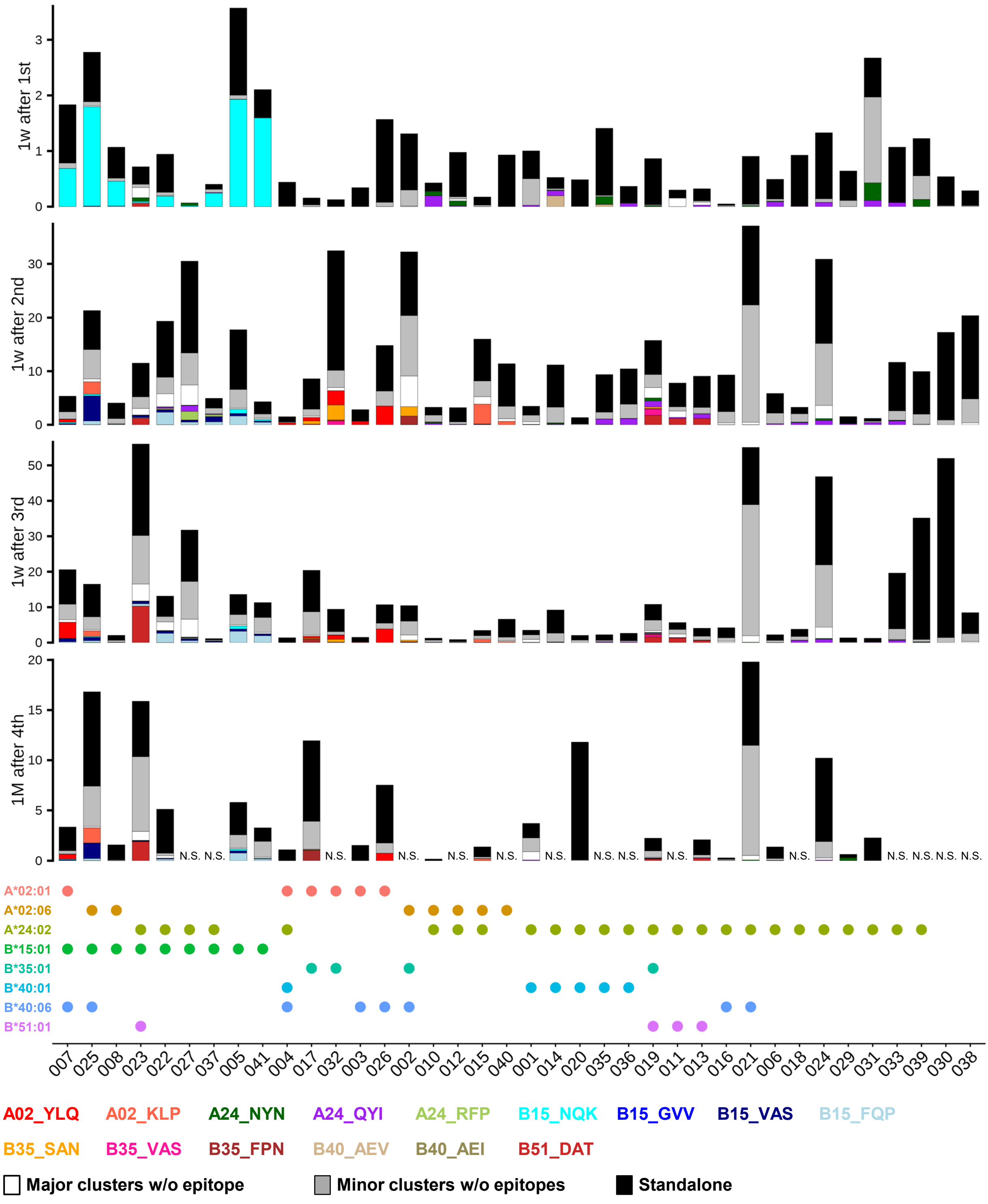

**Extended Data Fig. 6. Total size of T cell clones with or without predicted spike epitopes after each vaccination dose.**

X-axis represents individual donors. Clones without predicted epitopes are classified into the three groups: i) clones belonging to major $CD8^+$ TCR clusters (white), ii) clones belonging to minor $CD8^+$ clusters with at least two TCRs (grey), or iii) clones not connected to any other TCRs in the TCR similarity network ("*standalone*" TCRs, black). Donors' HLA alleles that present known spike epitopes are shown by colored circles below the bar plots. 1': 1 week after the $1^{st}$ dose. 2': 1 week after the $2^{nd}$ dose. 3': 1 week after the $3^{rd}$ dose. 4': 1 month after the $4^{th}$ dose. N.S.: Not sampled.

Extended Data Fig. 7

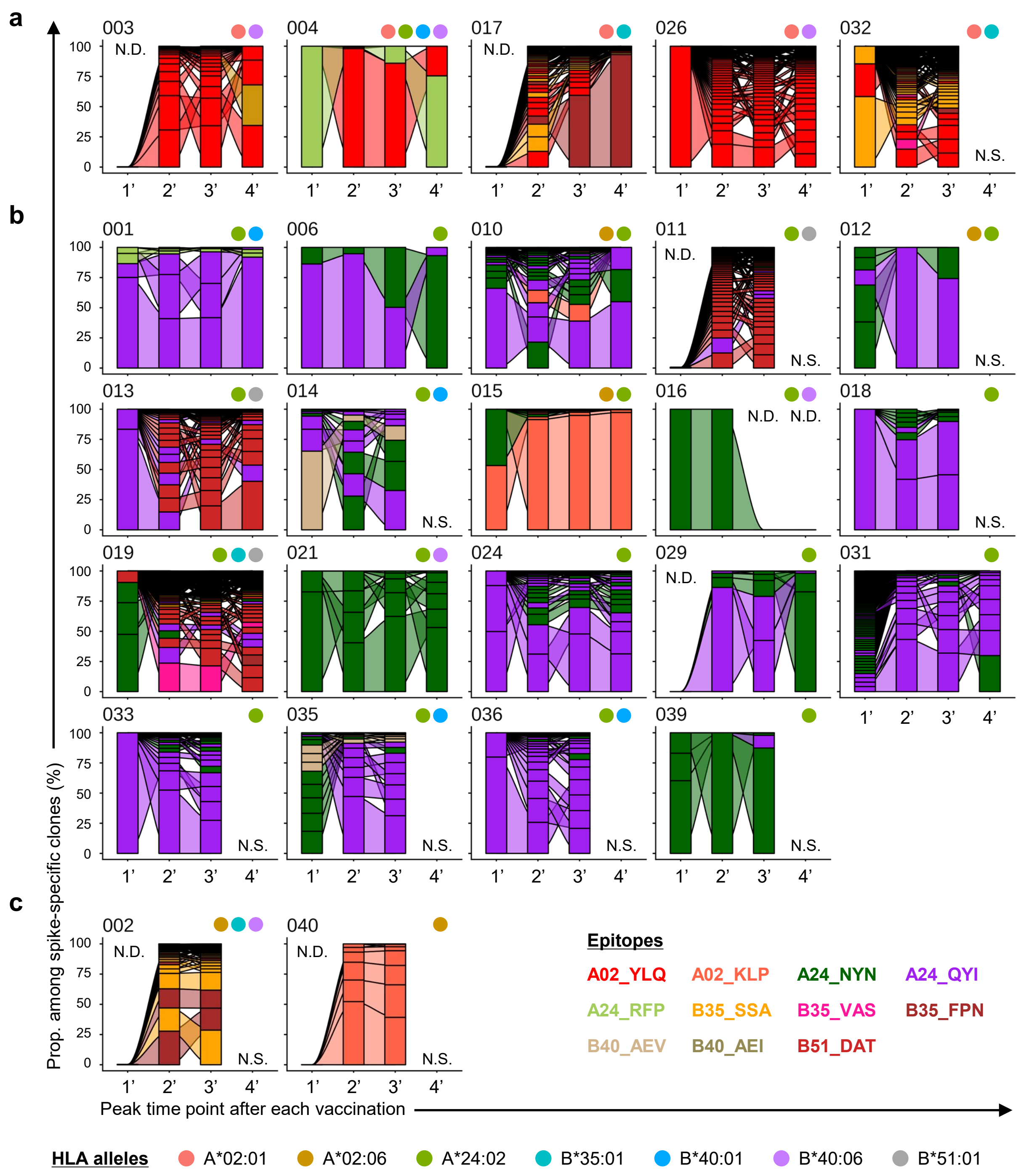

**Extended Data Fig. 7. Longitudinal kinetics of vaccine-associated clones de-orphanized to spike epitopes in B*15:01$^{-}$ donors.**

(**a**-**c**) Each bar, connected by ribbons, represents an individual CD8$^{+}$ T cell clone de-orphanized to a spike epitope. Bar height represents the proportion of each clone among those de-orphanized to spike epitopes, and bar colors correspond to respective epitopes. Numbers and colored circles above the plots denote individual donors and HLA alleles they carry. Only HLA alleles that present spike epitopes associated with TCR clusters (summarized in Fig. 4d) are shown. The plots of B*15:01$^{-}$ A*02:01$^{+}$ donors (**a**), B*15:01$^{-}$ A*02:01$^{-}$ A*24:02$^{+}$ donors (**b**), and the remaining donors (**c**) are shown. Donors 030 and 038 are excluded because they lack these HLA alleles, and Donor 020 is not shown due to the absence of de-orphanized CD8$^{+}$ clones. 1': 1 week after the 1$^{st}$ dose. 2': 1 week after the 2$^{nd}$ dose. 3': 1 week after the 3$^{rd}$ dose. 4': 1 month after the 4$^{th}$ dose. N.S.: Not sampled. N.D.: Not Detected.

Extended Data Fig. 8

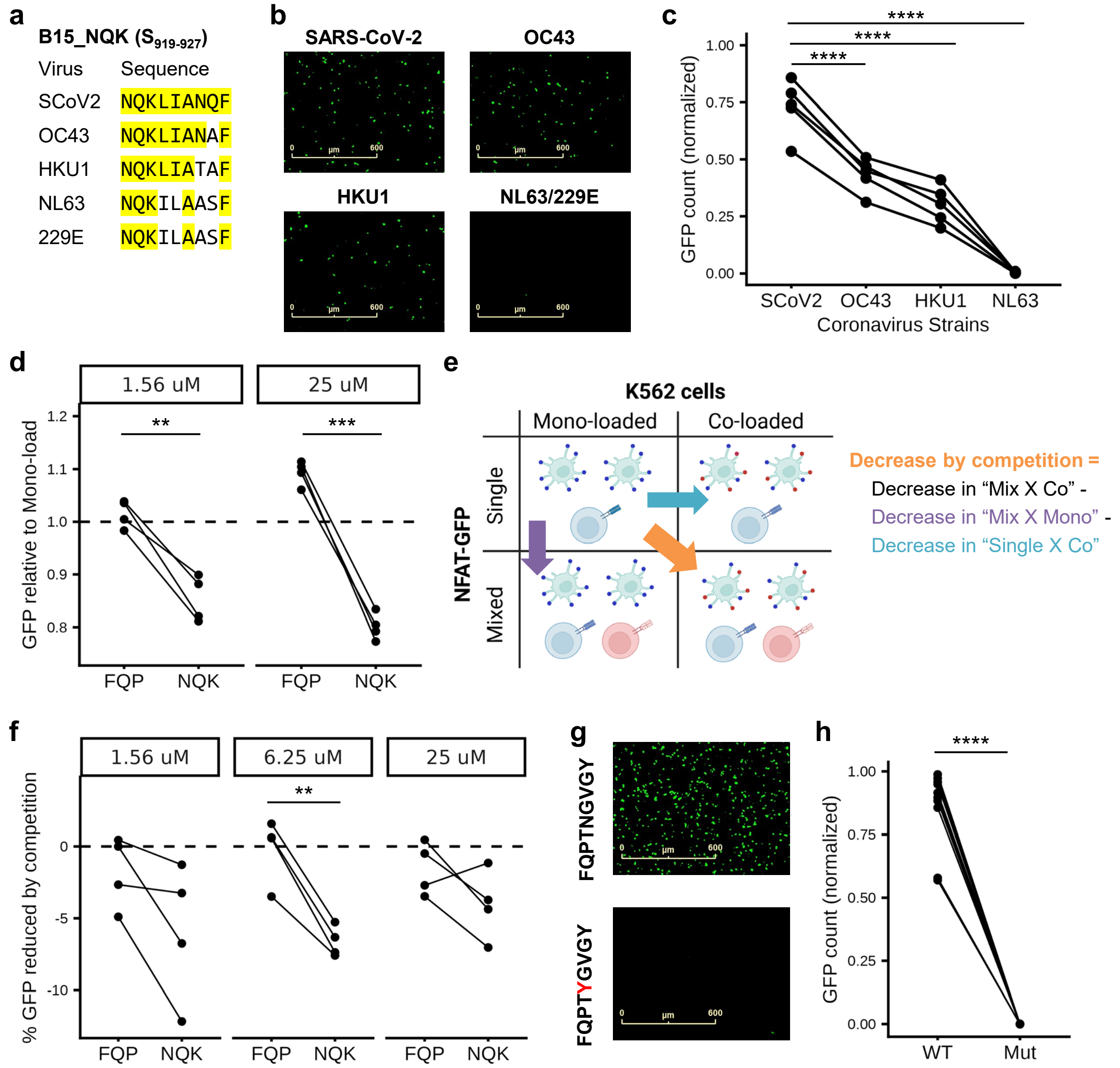

**Extended Data Fig. 8. Characterization of SARS-CoV-2 spike-specific TCRs targeting B*15:01-presented epitopes by in vitro co-culture assay.**
(**a**) The amino acid sequence of the B15_NQK epitope of SARS-CoV-2 and hCoVs (OC43, HKU1, and NL63/229E). Conserved amino acids are highlighted in yellow. (**b**) Representative Incucyte images showing GFP signals from NQK-specific TCR-transduced NFAT-GFP cells after co-culturing with HLA-B*15:01$^+$ K562 cells added with 50 μM NQK peptide of SARS-CoV-2 or hCoVs. (**c**) The count of GFP$^+$ NFAT-GFP reporter cells after co-culture. Data from the same TCR-transduced NFAT-GFP cells are connected with lines. BH-adjusted p-values of the paired Student's t-test were shown. (**d**) Competition effect between FQP- and NQK-specific TCRs, as quantified in Fig. 7e. 1.56 or 25 μM of NQK and/or FQP peptide is loaded on B15$^+$ K562 cells. (**e**) Schematic overview for evaluating the inter-TCR competition effect using *single vs mono-loaded* and *single vs co-loaded* conditions (see *Method*). (**f**) Inter-TCR competition effect measured as relative GFP reduction from the *single vs mono-loaded* condition. (**g**) Representative Incucyte images showing GFP signals from FQP-specific TCR-transduced NFAT-GFP cells after co-culturing with HLA-B*15:01$^+$ K562 cells added with 25 μM wild-type (top) or N5Y-mutated (bottom) FQP peptide. (**h**) The count of GFP$^+$ NFAT-GFP reporter cells after co-culture with wild-type ("WT") or mutated ("Mut") FQP peptide. P-values from the paired Student's t-test are shown (n=10). (**c**,**h**) The count of GFP$^+$ objects normalized to that of the positive control (co-cultured well with Cell Stimulation cocktail) is shown. (**d**,**f**) P-values from the paired Student's t-test are shown (n=4: 2 NQK-TCRs x 2 FQP-TCRs). ****$p<0.0001$; ***$p<0.001$; **$p<0.01$; *$p<0.05$. (**e**) Created using BioRender.com.

Supplementary Table 4

| Cluster | # Clones[1] | | Predicted Target (Summarized) | | SARS-CoV-2 spike-specific TCRs[2] | | HLA sharing across donors | | SARS-CoV-2-associated TCRβs[4] | | Other resources | | | In vitro validation[5] |
|---|---|---|---|---|---|---|---|---|---|---|---|---|---|---|
| | NMU | Ref. | **HLA** | **Peptide** | HLA | Peptide | HLA | # Donors[3] | **HLA** | **Peptide** | **HLA** | **Peptide** | **Resource** | |
| 1 | 154 | 0 | B*15:01 | | | | B*15:01 | 8 / 10 | | . | | | | |
| 2 | 127 | 0 | **B*51:01** | **DATNVVIKV** | | | A*24:02 | 10 / 14 | **B*51:01** | . | **B*51:01** | **DATNVVIKV** | Adragna et al. | 5 / 7* |
| 3 | 98 | 8 | **A*24:02** | **NYNYLYRLF** | **A*24** | **NYNYLYRLF** | A*24:02 | 16 / 16 | A*24:02 | NYNYLYRLF | | | | |
| 4 | 93 | 0 | **B*35:01** | **SSANNCTFEY** | | | B*35:01 | 4 / 5 | **B*35:01** | **SSANNCTFEY** | | | | 4 / 4 |
| 5 | 65 | 6 | **A*24:02** | **NYNYLYRLF** | **A*24** | **NYNYLYRLF** | A*24:02 | 17 / 17 | A*24:02 | NYNYLYRLF | A*24:02 | NYNYLYRLF | Structural modeling | 4 / 4 |
| 6 | 47 | 8 | **A*02:01** | **YLQPRTFLL** | **A*02** | **YLQPRTFLL** | A*02:01 | 6 / 7 | A*02:01 | YLQPRTFLL | A*02:01 | YLQPRTFLL | Structural modeling | 10 / 10 |
| 7 | 49 | 0 | **A*24:02** | **RFPNITNLCPF** | | | A*24:02 | 6 / 9 | **A*24:02** | **RFPNITNLCPF** | | | | |
| 8 | 48 | 0 | A*26:01 | | | | A*26:01<br>*A*24:02* | 4 / 7 | **A*26:01** | . | | | | |
| 9 | 43 | 3 | **A*02:01** | **YLQPRTFLL** | **A*02** | **YLQPRTFLL** | A*02:01 | 5 / 6 | A*02:01 | YLQPRTFLL | A*02:01 | YLQPRTFLL | Structural modeling | 10 / 10 |
| 10 | 44 | 0 | **B*51:01** | **DATNVVIKV** | | | *A*24:02*<br>B*51:01 | 4 / 5 | **B*51:01** | . | **B*51:01** | **DATNVVIKV** | Adragna et al. | 3 / 3 |
| 11 | 43 | 0 | **B*15:01** | **FQPTNGVGY** | | | B*15:01 | 9 / 9 | | . | **B*15:01** | **FQPTNGVGY** | Predicted strong binders | 5 / 5 |
| 12 | 40 | 2 | **A*24:02** | **NYNYLYRLF** | **A*24** | **NYNYLYRLF** | A*24:02 | 12 / 12 | A*24:02 | NYNYLYRLF | | | | |
| 13 | 34 | 0 | **B*15:01** | **VASQSIIAY** | | | **B*15:01** | 8 / 8 | | **VASQSIIAY**<br>**SQSIIAYTM** | B*15:01 | VASQSIIAY | Structural modeling | 5 / 5 |
| 14 | 33 | 0 | **B*51:01** | **DATNVVIKV** | | | *A*24:02*<br>B*51:01 | 4 / 4 | **B*51:01** | . | **B*51:01** | **DATNVVIKV** | Adragna et al. | 2 / 4* |
| 15 | 32 | 0 | **B*15:01** | **FQPTNGVGY** | | | B*15:01 | 8 / 8 | | | **B*15:01** | **FQPTNGVGY** | Structural modeling | 5 / 5 |
| 16 | 29 | 2 | **B*15:01** | **NQKLIANQF** | **B*15** | **NQKLIANQF** | B*15:01 | 8 / 8 | B*15:01 | NQKLIANQF | | | | 5 / 5 |
| 17 | 30 | 0 | **A*24:02** | **QYIKWPWYI** | **A*24** | **QYIKWPWYI** | A*24:02 | 15 / 15 | | . | | | | 5 / 5 |
| 18 | 30 | 0 | B*44:03 | | | | *A*33:03*<br>B*44:03 | 4 / 4 | B*44:03 | . | | | | |
| 19 | 27 | 0 | B*54:01 | | | | *A*24:02*<br>B*54:01 | 5 / 6 | | . | | | | |
| 20 | 27 | 0 | B*35:01 | | | | B*35:01 | 4 / 4 | B*35:01<br>*C*04:01* | . | | | | |
| 21 | 24 | 0 | | | | | B*35:01 | 3 / 3 | A*24:02 | | | | | |
| 22 | 23 | 0 | B*55:02 | | | | *A*24:02*<br>B*55:02 | 4 / 6 | | . | | | | |
| 23 | 23 | 0 | B*15:01 | | | | B*15:01 | 8 / 8 | | . | | | | |
| 24 | 22 | 0 | **B*15:01** | **VASQSIIAY** | | | **B*15:01** | 8 / 8 | | **VASQSIIAY**<br>**SQSIIAYTM** | **B*15:01** | **VASQSIIAY** | Structural modeling | |
| 25 | 21 | 0 | **A*24:02** | **QYIKWPWYI** | **A*24** | **QYIKWPWYI** | A*24:02 | 12 / 12 | | . | | | | 5 / 5 |

**Supplementary Table 4. Predicted peptide and HLAs for 25 major TCR clusters of CD8$^{+}$ vaccine-associated clones.** [1]The number of TCRs included in each cluster, which are derived from our dataset (*NMU*) or previous studies[24–26]. [2]Epitopes predicted by public databases of α/βTCRs such as VDJdb or identified in spike pHLA tetramer TCR-seq in our previous study[11,12]. [3]Number of donors with a corresponding HLA allele among those contributing CD8$^{+}$ clones to each cluster. [4]HLA allele and target epitope predicted by the most frequent edit-1 neighbor in the SARS-CoV-2-linked TCRβ datasets[34,35], which are shared across more than 20% of clones in each cluster. Only strong binder peptides predicted by NetMHCpan-4.1 are shown[33]. [5] Number of TCRs whose target epitope was validated by co-culture assay among those reconstituted on NFAT-GFP Jurkat cell line. *Co-culture assay was performed at 50 µM of the candidate peptide. Details of spike epitopes are summarized in Fig. 4d.

Supplementary Table 5

| Cluster | # Clones[1] | | Predicted Target (Summarized) | | SARS-CoV-2 spike-specific TCRs[2] | | HLA sharing across donors | | SARS-CoV-2-associated TCRβs[4] | |
|---|---|---|---|---|---|---|---|---|---|---|
| | NMU | Ref. | HLA | Peptide | HLA | Peptide | HLA | # Donors[3] | HLA | Peptide |
| 29 | 17 | 0 | A*02:06 | KLPDDFTGC | | | A*02:06 | 4 / 5 | | KLPDDFTGC |
| 30 | 17 | 0 | B*15:01 | VASQSIIAY<br>SQSIIAYTM | | | B*15:01 | 7 / 7 | | VASQSIIAY<br>SQSIIAYTM |
| 38 | 14 | 0 | A*02:01 | YLQPRTFLL | A*02 | YLQPRTFLL | A*02:01 | 5 / 5 | A*02:01 | YLQPRTFLL |
| 39 | 14 | 0 | B*15:01 | VASQSIIAY<br>SQSIIAYTM | | | B*15:01 | 4 / 4 | | VASQSIIAY<br>SQSIIAYTM |
| 47 | 12 | 0 | B*35:01 | SSANNCTFEY | | | B*35:01 | 4 / 4 | | SSANNCTFEY |
| 59 | 9 | 0 | B*35:01 | FPNITNLCPF<br>QPTESIVRF | | | B*35:01 | 3 / 3 | B*35:01 | FPNITNLCPF<br>QPTESIVRF |
| 62 | 9 | 0 | A*24:02 | QYIKWPWYI | A*24 | QYIKWPWYI | A*24:02 | 6 / 6 | A*24:02 | QYIKWPWYI |
| 66 | 8 | 0 | A*02:01 | YLQPRTFLL | A*02 | YLQPRTFLL | A*02:01 | 3 / 3 | A*02:01 | YLQPRTFLL |
| 67 | 8 | 0 | B*15:01 | GVVFLHVTY | | | B*15:01 | 5 / 5 | B*15:01 | GVVFLHVTY |
| 76 | 6 | 0 | A*24:02 | NYNYLYRLF | A*24 | NYNYLYRLF | A*24:02 | 5 / 5 | A*24:02 | NYNYLYRLF |
| 79 | 6 | 0 | B*15:01 | NQKLIANQF | B*15 | NQKLIANQF | B*15:01 | 5 / 5 | | NQKLIANQF |
| 85 | 6 | 0 | B*40:01 | AEVQIDRL | | | A*24:02<br>A*33:03<br>B*40:01 | 2 / 2 | B*40:01 | AEVQIDRL |
| 93 | 5 | 0 | A*24:02 | QYIKWPWYI | A*24 | QYIKWPWYI | A*24:02 | 5 / 5 | | . |
| 98 | 5 | 0 | A*24:02 | NYNYLYRLF | A*24 | NYNYLYRLF | A*24:02 | 4 / 4 | A*24:02 | NYNYLYRLF |
| 112 | 4 | 0 | B*35:01 | VASQSIIAY | | | B*35:01 | 2 / 2 | | VASQSIIAY |
| 113 | 4 | 0 | B*35:01 | FPNITNLCPF<br>QPTESIVRF | | | B*35:01 | 2 / 2 | B*35:01<br>C*04:01 | FPNITNLCPF<br>QPTESIVRF |
| 122 | 4 | 0 | B*15:01 | NQKLIANQF | B*15 | NQKLIANQF | B*15:01 | 2 / 2 | | NQKLIANQF |
| 123 | 4 | 0 | A*24:02 | NYNYLYRLF | | | A*24:02 | 2 / 2 | A*24:02 | NYNYLYRLF |
| 124 | 4 | 0 | A*24:02 | QYIKWPWYI | A*24 | QYIKWPWYI | A*24:02 | 3 / 3 | | . |
| 125 | 4 | 0 | B*15:01 | NQKLIANQF | B*15 | NQKLIANQF | B*15:01 | 3 / 3 | B*15:01 | NQKLIANQF |
| 129 | 4 | 0 | B*15:01 | NQKLIANQF | B*15 | NQKLIANQF | B*15:01 | 3 / 3 | | . |
| 131 | 4 | 0 | B*40:06 | AEIRASANL | | | B*40:06 | 3 / 3 | | AEIRASANL |

**Supplementary Table 5. Predicted peptide and HLAs for minor TCR clusters of $CD8^+$ vaccine-associated clones.** [1]The number of TCRs included in each cluster, which are derived from our dataset (*NMU*) or previous studies[24-26]. [2]Epitopes predicted by public databases of α/βTCRs such as VDJdb or identified in spike pHLA tetramer TCR-seq in our previous study[11,12]. [3]Number of donors with a corresponding HLA allele among those contributing $CD8^+$ clones to each cluster. [4]HLA allele and target epitope predicted by the most frequent edit-1 neighbor in the SARS-CoV-2 linked TCRβ datasets[34,35]. Only strong binder peptides predicted by NetMHCpan-4.1 are shown[35].

Supplementary Table 6

| Signature | Genes | No. genes |
|---|---|---|
| CD8_Naive_CCR7 | NT5E, CD8B, LEF1, FCER1G, CCR7, PASK, BACH2 | 7 |
| CD8_TCM_HAVCR2 | GZMK, HAVCR2, LGALS3, CD38, POU2AF1, CTLA4, TIGIT, CD27 | 8 |
| CD8_TEM_CMC1 | GZMK, DUSP2, EOMES, CCL5, CD160, TIGIT, CCL4, HLA-DQB1, CD74, CST7, GZMA, HLA-DPA1, CD8A, CXCR3, HLA-DRA, CD8B | 16 |
| CD8_TEM_GNLY | GNLY, GZMB, FCGR3A, GZMH, NKG7, CX3CR1, PRF1, KLRC3, ZNF683, CST7, CCL4, KLRF1, TBX21, CCL5 | 14 |
| CD8_TEM_ZNF683 | TRDC, ZNF683, IKZF2, KLRC3, CXCR3, IGHM, NCR3, CD79A, CD7, IFITM3 | 10 |
| IFNα (Msigdb) | CD74, CXCL10, CXCL11, IFITM2, IFITM3, IL15, IL4R, LAMP3, LAP3, SELL | 10 |

**Supplementary Table 6. List of genes used for calculating signature scores in Fig. 2k.** Among the marker genes for blood T cell subpopulations reported in Wang et al.[20] and the Hallmark IFNα response gene set from MSigDB[21], those included in the BD Rhapsody Immune Response Panel are listed.
